# RNA binding by ADARs prevents RNA interference from attacking self-produced dsRNA

**DOI:** 10.1101/2025.08.28.672520

**Authors:** Nabeel S. Ganem, Dean Light, Roni Haas, Boaz Goldstein, Alla Fishman, Yahav Festinger, Noa Ben-Asher, Orna Ben-Naim Zgayer, Berta Eliad, Fabian Glaser, Suba Rajendren, Heather A. Hundley, Ayelet T. Lamm

## Abstract

The ability of an organism to identify self and foreign RNA is central to eliciting an immune response in times of need while avoiding autoimmunity. As viral pathogens typically employ double-stranded RNA (dsRNA), host identification, modulation, and response to dsRNA is key. However, dsRNA is also abundant in host transcriptomes, raising the question of how these molecules can be differentiated. Two host pathways that regulate dsRNA are A-to-I RNA editing by adenosine deaminases (ADARs), and RNA interference (RNAi). Both mechanisms are important for normal organism development and function by regulating gene expression. Herein, we studied the structure and amount of siRNAs at editing sites and the ability of ADARs to prevent exogenous RNAi using the model organism, *Caenorhabditis elegans*. We found that the number of siRNAs targeting edited genes is significantly upregulated in ADAR mutant animals. We also found that despite an almost complete depletion of primary siRNAs generated from editing sites in wildtype animals, secondary siRNAs are generated from edited transcripts, suggesting ADARs antagonize only the first step of RNAi processing. We show that ADARs interfere with the efficacy of exogenous RNAi *in vivo*, probably to prevent trans-silencing, and have indications that ADR-2 binding to the dsRNA is needed for the efficient prevention of RNAi. This work sheds light on how the RNA editing process protects self-produced dsRNAs from aberrant recognition by the immune processes in the cell and from by-product degradation.

**Graphical abstract:** 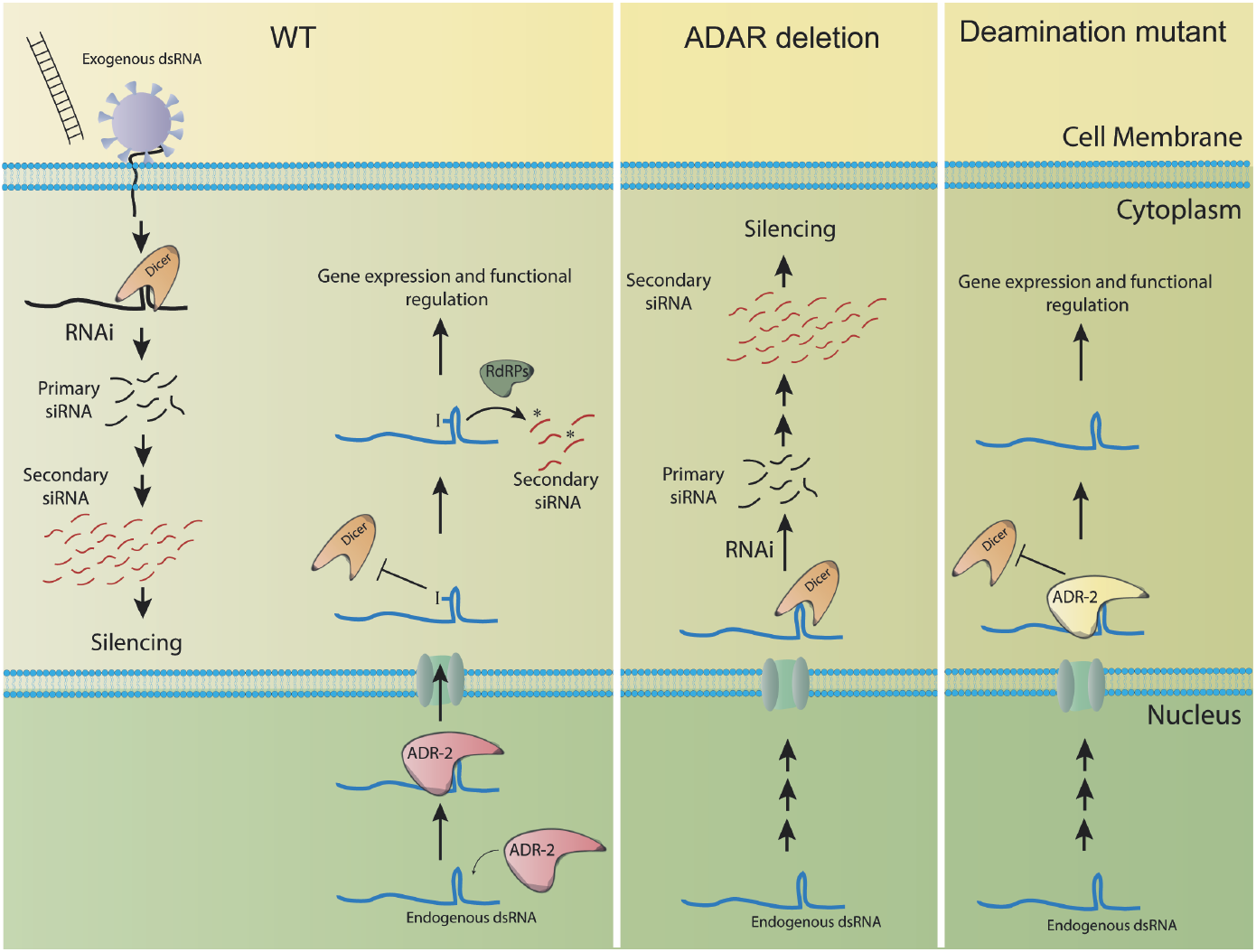

## Introduction

A-to-I RNA editing is one of the most abundant RNA modifications in the human transcriptome and is common in metazoans (Bajad et al., 2017; Schaffer and Levanon, 2021; Song et al., 2022; Zhang et al., 2023). The editing process occurs specifically on dsRNAs and is catalyzed by Adenosine deaminases that act on RNA (ADARs) enzymes (Bass, 2002). Editing sites mainly reside in repetitive genomic structures, in introns, and within the 3’ untranslated region (UTR) of genes (Athanasiadis et al. 2004; Blow et al. 2004; Kim et al. 2004; Levanon et al. 2004; Levanon et al. 2005; Barak et al. 2009; Li et al. 2009; Kleinberger and Eisenberg 2010; Osenberg et al. 2010; Paz-Yaacov et al. 2010). The scope of the developmental and regulatory role of RNA editing is just beginning to emerge. However, it is well established that RNA editing is essential in mammals, and altered editing patterns in humans and mice have been linked to autoimmune diseases, neuropathological disorders, and various tumors (Erdmann et al., 2021). Studies in mammals suggested a role for RNA editing to protect against wrongly triggering the immune response by the interferon when encountering self-produced dsRNAs in the cytoplasm (George et al., 2016; Liddicoat et al., 2015; Mannion et al., 2014). ADARs affect the interferon reaction both by editing and by binding to the dsRNA (reviewed in (Cottrell et al., 2024; Hu and Li, 2024; Liddicoat et al., 2016)).

While not essential, disrupting the editing pathways in *C. elegans* and *Drosophila* results in behavioral and neural defects (Ganem et al., 2019; Hartner et al., 2004; Hartner et al., 2009; Jepson and Reenan, 2009, 2010; Jin et al., 2009; Sebastiani et al., 2009; Tonkin et al., 2002; Wang et al., 2000; Wang et al., 2004). In *C. elegans*, there are two ADAR genes, *adr-1 and adr-2* (Tonkin et al., 2002). ADR-2 is the only active enzyme; however both genes are part of the editing process (Arribere et al., 2020). ADR-1 was shown to bind dsRNA and to regulate RNA editing efficiency of specific adenosines by promoting the interaction of ADR-2 with substrates (Ganem et al., 2019; Rajendren et al., 2018; Washburn et al., 2014). Mutations in either *adr-1* or *adr-2* are not lethal but cause developmental defects, including chemotaxis defects, changes in the lifespan, egg laying defects, and others (Dhakal et al., 2024; Ganem et al., 2019; Mahapatra et al., 2023; Reich et al., 2018; Sebastiani et al., 2009; Tonkin et al., 2002). In *C. elegans*, RNA editing is thought to have an antagonistic effect on RNAi. However, how ADARs interfere with RNAi processing and at which stage is not yet known. Indications come from transcriptomics studies that showed enrichment of siRNAs in hyper-edited genomic regions in the absence of ADARs (Goldstein et al., 2017; Reich et al., 2018; Warf et al., 2012; Wu et al., 2011). In addition, some of the phenotypes observed in ADAR mutant worms are rescued by introducing another mutation in RNAi machinery factors (*rde-1* and *rde-4*) (Tonkin and Bass, 2003). Similarly, introducing mutations in *rde-1* and *rde-4* in ADAR mutant worms rescued lncRNA downregulation at the embryonic developmental stage (Goldstein et al., 2017). Studies using *C. elegans* laid the groundwork for the study of RNAi in many different organisms (Fire, 2007). Most RNAi processes in *C. elegans* generate two populations of small RNAs (Almeida et al., 2019). The first population contains primary siRNAs, which are cleaved from dsRNA by Dicer. They can be 26nt siRNA products of endogenous RNAi or 23nt siRNAs products of viral infection. Recently, endogenous 23H-RNAs were also identified (Knittel et al., 2024). The second population contains secondary 21-22nt siRNA, which are polymerized by RNA-dependent-RNA-polymerase (RdRPs) (Grishok, 2013). It was shown that genes in proximity to editing-enriched regions (EERs), pseudogenes, and lncRNAs are downregulated in ADAR mutants, and that this downregulation is dependent on RNAi (Goldstein et al., 2017; Reich et al., 2018). RRF-3 is an RdRP protein required for 26G endo-siRNAs biogenesis (Gent et al., 2010). Worms harboring a mutation in *rrf-3* gene exhibit hypersensitivity and stronger silencing by the RNAi machinery (Simmer et al., 2002). RRF-3 mutation, together with mutations in ADAR genes, causes developmental abnormalities (Reich et al., 2018). These phenotypes were rescued by introducing additional mutations in other antiviral RNAi factors, such as DRH-1 (which is involved in RNAi processing of viral RNA), RDE-1 (which is needed for exogenous RNAi), and others (Reich et al., 2018). In another study, both ADAR genes and the eri-6\7 pathway were shown to regulate the silencing of LTR retrotransposons and endogenous viral elements (Fischer and Ruvkun, 2020). Together, these studies suggest that RNA editing by ADARs in *C. elegans* protects dsRNA from RNAi, probably to prevent an aberrant immune response by RNAi against self-produced dsRNAs. However, how ADARs interfere with RNAi processing and at which stage is not yet known.

Possible hypotheses on how RNA editing prevents RNAi in *C. elegans* are that accumulation of inosine prevents Dicer from binding the dsRNA, that A-to-I nucleotide changes can also change the structure of the dsRNA, preventing efficient Dicer binding, or that ADR-1 and ADR-2 compete with Dicer for binding the dsRNA (reviewed in (Ganem and Lamm, 2017)). ADR-2 was shown to have editing-independent functions, for example, it regulates the switch between spermatogenesis and oogenesis (Erdmann et al., 2024), possibly through its RNA binding activity. Additionally, the subcellular localization of ADR-2 may affect its function. ADR-2 nuclear localization is regulated by ADBP-1 protein (Eliad et al., 2024; Ota et al., 2013). However, ADR-2, even in the cytoplasm, can edit dsRNAs but to a lesser extent (Eliad et al., 2024; Mu et al., 2025). To further understand the relationship between RNA editing and RNAi, we analyzed the amount and structure of siRNAs generated from editing sites in non-repetitive regions and specifically from the 3’UTRs of genes. In addition, we studied whether exogenous RNAi is also affected by RNA editing and how RNA editing interferes with RNAi.

## Results

### Primary and secondary siRNAs are upregulated at 3’UTR edited genes in ADAR mutant worms

Several studies showed that RNA editing prevents the processing of dsRNAs by RNAi in *C. elegans* (Fischer and Ruvkun, 2020; Goldstein et al., 2017; Reich et al., 2018; Warf et al., 2012; Wu et al., 2011). In *C. elegans*, RNAi is the main immunity process. However, RNAi is also used endogenously in the worm to silence genes. It is unclear how RNA editing affects RNAi, and it is possible that the two steps of RNAi (i.e. 1. Recognition and generation of primary siRNAs and 2. Amplification and generation of secondary siRNAs by RdRPs) may be affected differentially by RNA editing. To better understand how RNA editing antagonizes RNAi, we sought a comprehensive examination of the siRNA repertoire in cells. Therefore, we generated siRNA libraries using the QsRNA-seq method (Fishman et al., 2018), capturing mainly primary siRNAs or secondary siRNAs from wildtype and ADAR mutant worms at the embryo stage, in which editing is strongest (Zhao et al., 2015). To make the analysis stringent, we concentrated on editing sites in non-repetitive regions that we identified in previous work, and specifically on editing sites in the 3’UTR of genes (Ganem et al., 2019; Goldstein et al., 2017). We concentrated on 3’UTR edited genes because, as in (Goldstein et al., 2017), we wanted to use sites that are uniquely related to genes and have high editing levels, while the number of sites in exons is too low for meaningful analysis (Goldstein et al., 2017). In addition, we used two ADAR mutant strains with different deletion alleles (*adr-1 (gv6)*; *adr-2 (gv42)* and *adr-1 (tm668)*; *adr-2 (ok735)*; see materials and methods).

Firstly, by comparing the number of siRNAs aligned to each gene in wildtype and ADAR mutants (Supp Table 1), we prepared a list of genes that have at least two-fold more siRNAs in ADAR mutants than in wildtype with Padj value < 0.05 for each method of siRNAs preparation, primary or secondary. To test if genes with enriched siRNAs in ADAR mutants compared to wildtype are downregulated in the ADAR mutants at the mRNA levels, we tested their expression using RNA-seq libraries generated in parallel to the siRNA libraries and published in (Goldstein et al., 2017). We did not observe significant downregulation when testing genes with enriched primary siRNAs in both ADAR mutant worms compared to all genes (Figure 1A, Supp Figure 1A, two-tailed T-test p-value non-significant). However, a mild downregulation was observed for genes enriched with secondary siRNAs in both ADAR mutant worms compared to all genes (Figure 1A P-value < e-7; Supp Figure 1A P-value < 0.01). Analyzing only 3’UTR edited genes enriched with siRNAs in ADAR mutants, we found that these genes are significantly downregulated at the mRNA level (Figure 1B and Supp Figure 1B, p-value < 0.01). This result is consistent with the downregulated expression of 3’UTR edited genes observed in ADAR mutants (Goldstein et al., 2017). Comparing counts per gene of primary siRNAs in wildtype and ADAR mutant worms, we observed that 3’UTR edited genes have a significant upregulation in the number of siRNAs in ADAR mutant worms (Figure 1C, Supp Figure 1C p-value < e-6). However, for the secondary siRNAs, this upregulation was only observed for one of ADAR mutants, *adr-1 (tm668)*; *adr-2 (ok735)* (Figure 1D, P-value < 0.003, Supp Figure 1D, P-value non-significant). In addition, 3’UTR edited genes had a very low number of primary siRNAs in wildtype worms (Figure 1C, Supp Figure 1C). We concluded that in the absence of RNA editing, 3’UTR edited genes are downregulated and accumulate both primary and secondary siRNAs, while the overall genes with siRNA accumulation in ADAR mutant worms are not significantly downregulated. This downregulation and the accumulation of siRNAs were observed before in several studies (Goldstein et al., 2017; Reich et al., 2018; Warf et al., 2012; Wu et al., 2011), suggesting a tight connection between expression downregulation and siRNA accumulation.

**Figure 1.**
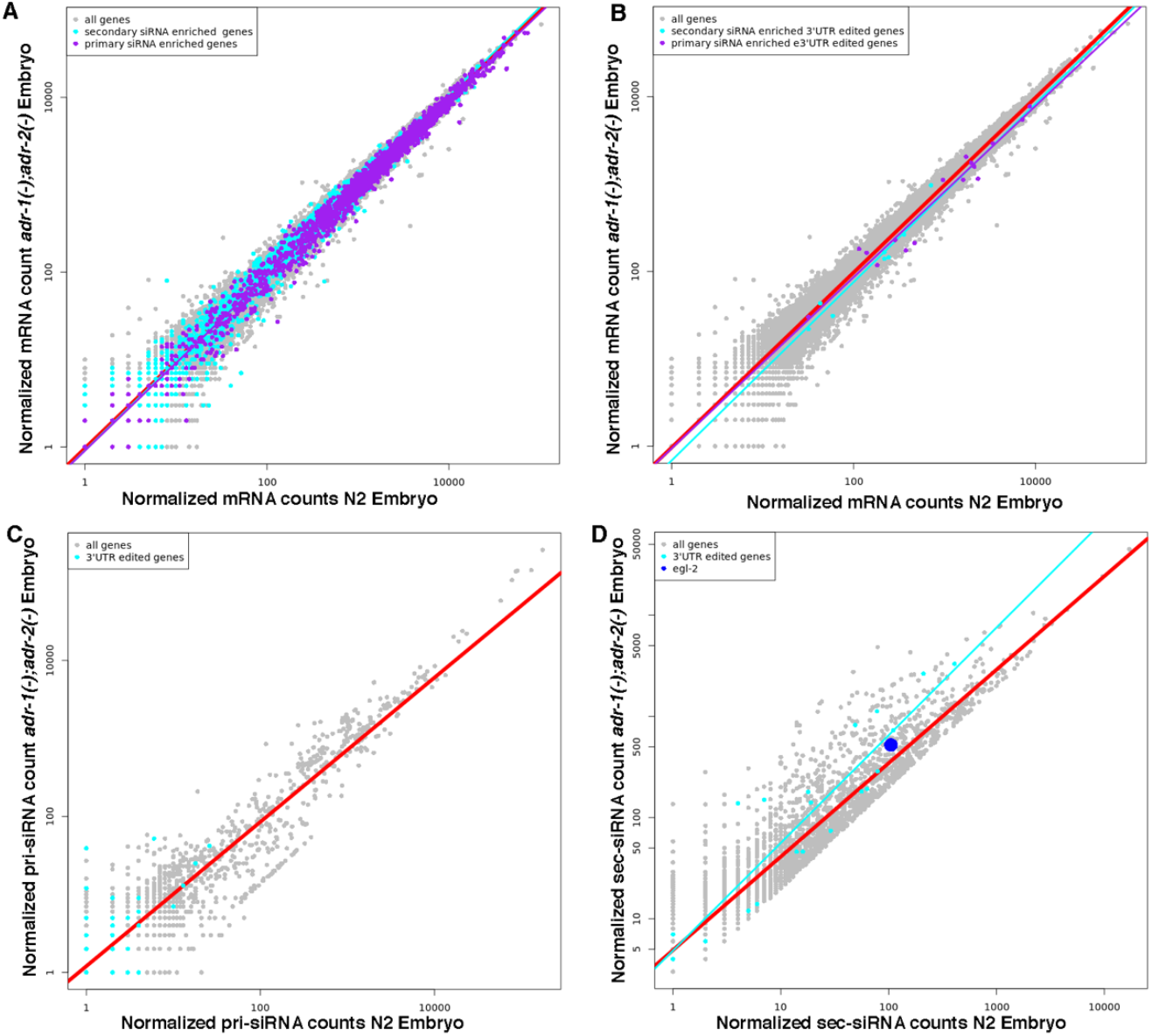
Genes edited at their 3’UTR are siRNAs enriched in ADAR mutant worms. Log scale plots presenting normalized mRNA sequence counts (A,B), primary siRNAs sequence counts (C), or secondary siRNAs sequence counts (D) of genes from at least 3 biological replicas in wildtype (N2) worms compared to ADAR mutant worms (*adr-1 (tm668)*; *adr-2 (ok735)*). Every dot in the graphs represents a gene. The red line is the regression line for all genes. Genes enriched in secondary siRNAs in ADAR mutant worms are marked in cyan (A,B) and their regression line is presented in cyan. Genes enriched in primary siRNAs in ADAR mutant worms are marked in purple (A,B) and their regression line is presented in purple. 3’UTR edited genes are marked in cyan (C,D) their regression line is presented in cyan. *egl-2* gene is marked in blue (D).

### Primary siRNAs are not generated from edited sites

We further examined the structure and size of siRNAs generated from editing sites. For this analysis, we only took small RNA sequences with antisense alignment to genes to observe the total size distribution of siRNAs. For the editing sites, we concentrated on 3’UTR editing sites in non-repetitive regions because they are well annotated (Ganem et al., 2019). For both primary and secondary siRNAs there was a significant enrichment of 23-mer siRNAs in both ADAR mutant worms compared to the wildtype worms at editing sites (Figure 2A,B, P-value<0.05 by one-tail T-test). A similar trend was observed for all sequences aligned to the transcriptome for the primary siRNAs but not for the secondary siRNAs (Figure 2C,D). The size distribution of primary and secondary siRNAs in the entire transcriptome was as expected, reflecting the different RNAi pathways e.g., for primary siRNAs 26-mers for endogenous RNAi, 22,23-mer exogenous and viral RNAi, and 21-mer for 21u, and for the secondary siRNAs the majority were RdRPs products e.g., 22-mers. However, at editing sites for primary siRNAs, most sequences were 23-mers, mainly reflecting the exogenous and viral RNAi pathways (Figure 2A compared to Figure 2C). This observation was also shown before by (Knittel et al., 2024; Reich et al., 2018; Wu et al., 2011), suggesting that RNA editing mainly antagonizes the viral RNAi pathway. To determine if the enrichment of siRNAs in ADAR mutants at editing sites is specific to 23-mer siRNAs, we took primary antisense siRNAs aligned to genes, sizes 23-mer, 25-mer, and 26-mer. We chose 26-mer to compare endogenous siRNAs to 23-mer viral siRNAs, and 25-mer as a control for siRNAs not associated with a certain pathway. For this analysis, we combined all primary siRNAs databases for each strain, e.g., all biological replicas. By comparing all genes with siRNAs alignment to edited genes from (Goldstein et al., 2017) with the different sizes, we found that 23-mer siRNAs are more aligned to edited genes than 25-mer or 26-mer siRNAs (Supp Figure 2). In addition, by comparing the alignment to edited genes in wildtype versus ADAR mutants, we found that only 23-mer siRNAs are significantly enriched (Supp Figure 3). Thus, our results suggest that although siRNAs of all sizes are aligned to edited genes, RNA editing inhibition of RNAi is specific to generating 23-mers, viral siRNAs.

**Figure 2.**
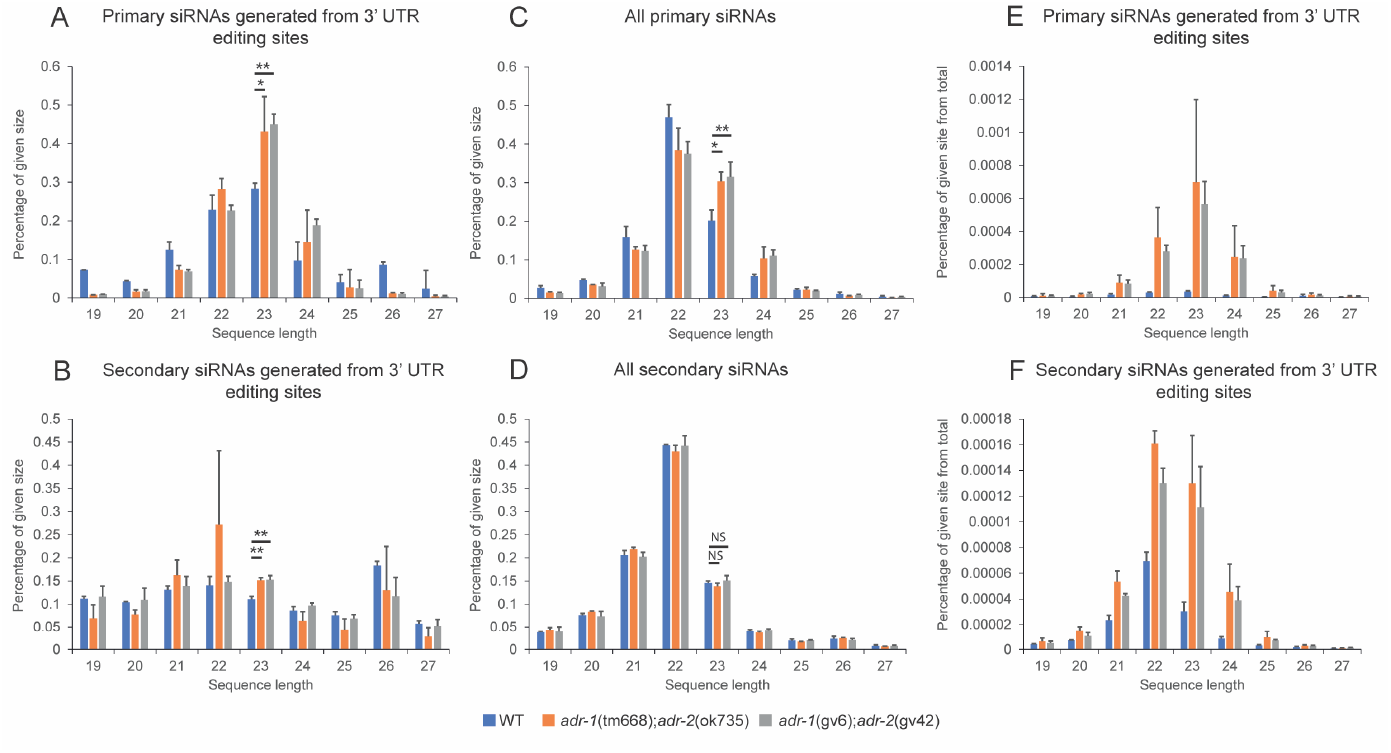
siRNAs are significantly depleted from editing sites in wildtype worms. Bar plot representing the percentage of antisense siRNA reads in a certain sequence length out of all reads of primary siRNA (A,C) or secondary siRNAs (B,D) aligned to 3’UTR non-repetitive editing sites (A,B) or to all sites (C,D). * p-value < 0.05, ** p-value < 0.01, calculated by one-tail T-test. (E,F) Bar plots showing the percentage of reads overlapping editing sites out of all reads, stratified by read length (19nt-27nt). Reads from enriched primary siRNA sequencing are in (E), and reads from enriched secondary siRNA sequencing are in (F). Wild type counts are shown in blue and ADAR mutants are shown in red (*adr-1 (tm668)*; *adr-2 (ok735)*) and grey (*adr-1 (gv6)*; *adr-2 (gv42)*). siRNAs in each sample were estimated independently and stdv is presented.

**Figure 3.**
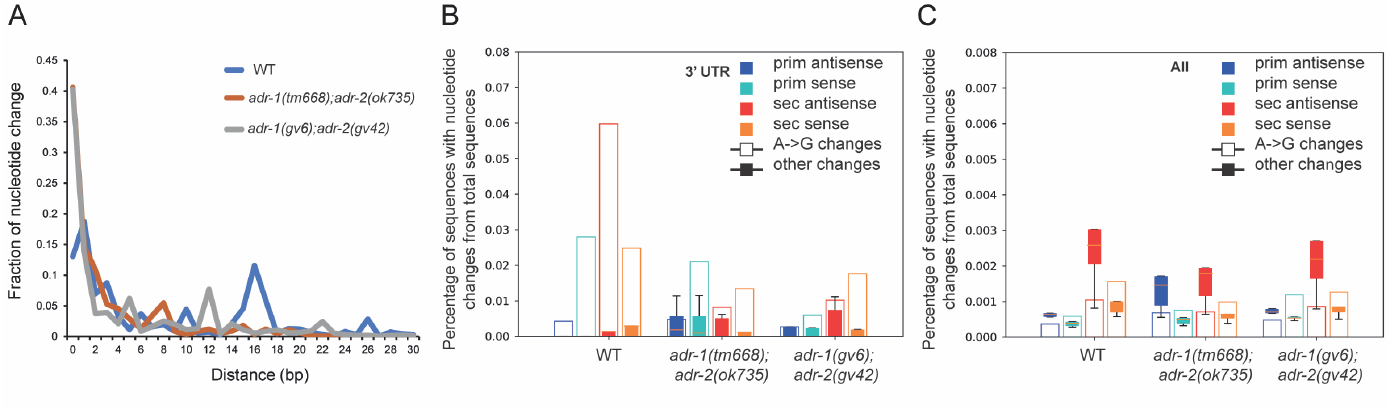
Secondary siRNAs aligned to editing sites contain ADAR editing changes. A. Distribution of distance between secondary siRNAs and the nearest primary siRNA found. X axis is distance in nucleotides and Y axis is the frequency of the distance in percentage. Wildtype, *adr1(tm668); adr-2(ok735)) and adr-1(gv6); adr-2(gv42))* are shown in blue, red and grey, respectively. (B,C) Distribution of percentage of editing events by editing type, stratified by sample type (primary or secondary, sense or anti-sense alignment) and genotype. In each sub-axis, open bars mark the A-to-G change, and filled bars represent the distribution of all other types of editing changes. (B) sites aligned to 3’UTR editing sites. (C) all sites.

We next compared the number of siRNAs in editing sites normalized to the number of siRNAs aligned to the entire transcriptome in wildtype or ADAR mutants based on the siRNAs size. We found a significant depletion of primary siRNAs in editing sites of all siRNA sizes in wildtype worms compared to ADAR mutant worms (Figure 2E, p-value < 0.05 for sizes 21-24). We also observed a significant depletion in the number of secondary siRNAs in wildtype worms (Figure 2F, p-value < 0.05 for sizes 21-24), but while there are secondary siRNAs in wildtype worms, most of the 3’UTR edited genes do not have any primary siRNAs aligned to edited sites at all in wildtype worms.

Primary siRNAs together with Argonautes recruit RdRPs to generate secondary siRNAs, which cover sequences mostly downstream but also upstream to the dsRNA trigger (Pak et al., 2012). Because most of the editing sites are in repetitive regions in the genome, it is possible that primary siRNAs that were generated in other non-edited locations in the genome and are similar in sequence to editing sites, can still serve to recruit RdRPs to generate a small number of secondary siRNAs. To test this theory, we analyzed the distance between uniquely aligned antisense secondary siRNAs to 3’UTR editing sites and non-uniquely aligned antisense primary siRNAs in wildtype worms and ADAR mutant worms. In both ADAR mutant worms, as expected, the distance between primary siRNAs and secondary siRNAs was very short, in most cases, they were adjacent (40% of the sequences have 0 nt distance, Figure 3A). In wildtype worms, the distance between primary and secondary siRNAs was significantly different from ADAR mutant worms (p-value < e-171, calculated using Anderson-Darling test (Scholz and Stephens, 1987)). In wildtype, many of the secondary siRNAs were far apart from the closest primary siRNAs, and only about 13% of the secondary siRNA sequences were adjacent to the primary siRNAs (0 nt distance).

We concluded that primary siRNAs are not generated from RNA that undergoes editing, although secondary siRNAs can be generated, probably using primary siRNAs that were generated from different sites, which are similar in sequence.

### Edited mRNA can serve as a template for secondary siRNAs generation

Although we almost detected no primary siRNAs generated from editing sites in wildtype worms, we still observed secondary siRNAs, though their number was significantly reduced. This result raised the question of whether siRNAs normally generated from editing loci bear the editing changes. To answer this question, we used the same editing sites at the 3’UTR of genes and performed an analysis to test whether siRNAs generated from these sites also have the A-to-G change. We compared nucleotide change frequencies for all nucleotide changes or specifically A-to-G changes in wildtype and ADAR mutants on editing sites in 3’UTR (Figure 3B) or on the entire transcriptome (Figure 3C). Interestingly, antisense secondary siRNAs generated from editing sites were enriched to the A-to-G change (Figure 3B). The percentage of secondary siRNAs containing editing changes was about 6% for all the 3’UTR edited genes, which is significant compared to other changes (P-value= 1.3e-6, calculated by one-tailed T-test). We presume that the percentage of edited secondary siRNA is an underestimation, as being short, siRNA with multiple editing-derived mismatches would fail to align and thus would be excluded from the analysis. While the number of secondary siRNAs was low in wildtype worms compared to ADAR mutant worms, it was enough for the statistical analysis. The number of primary siRNAs was too low for an unequivocal conclusion. When analyzing primary and secondary siRNAs that aligned to the entire transcriptome, we did not observe enrichment of siRNAs containing the editing change (Figure 3C) as was observed in editing sites. Enrichment of secondary siRNAs with the editing change was also observed when examining specific genes (an example can be seen in Supp Figure 4). This result, together with our previous results, suggests that if the RNA editing pathway successfully targets a dsRNA, it will not be processed by the first step of the siRNA pathway, i.e., cleavage by Dicer. This results in fewer primary siRNAs targeting editing sites, and, in turn, fewer secondary siRNAs are synthesized over the transcript. Secondary siRNAs bearing T-to-C changes at editing sites indicate that edited transcripts can still be used as a template for secondary siRNA synthesis, by RdRPs (guided by primary siRNA produced from unedited dsRNA). Thus, RNA editing only interferes with the generation of primary siRNAs and does not affect the generation of secondary siRNAs by RdRPs.

**Figure 4.**
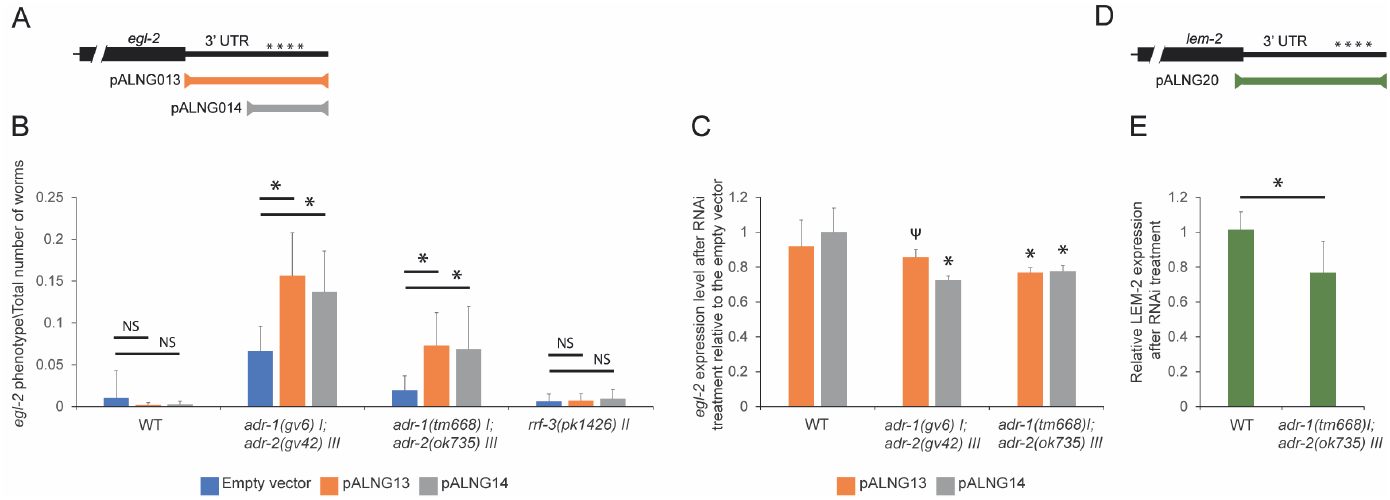
RNA editing at genes 3’ UTR prevents RNAi efficiency. (A) A scheme showing RNAi vectors targeting *egl-2* 3’ UTR; pALNG013 (323 bp) and pALNG014 (182 bp). Asterisks indicate editing sites. (B) Quantification of the fraction of *egl-2* phenotype from total worms observed in the RNAi experiments. Abnormal phenotypes that were scored include bloated worms, exploded worms, egg retention, and bag of worms phenotypes. P-values were calculated from at least 3 biological replicas, and the stdv is shown; NS for non-significant, *P-value < 0.05. The mutant strain, *rrf-3*, was used to control for RNAi hypersensitivity (Simmer et al., 2002). (C) Real-time PCR on RNA produced from worms after RNAi treatment. The empty RNAi plasmid was used for control. The experiment was repeated 3 times. Statistical analysis was done by comparing mutant strains with wildtype under the same RNAi treatment. * P-value < 0.05; ^Ψ^ P-value = 0.055103. (D) A scheme showing RNAi vector targeting *lem-2* 3’ UTR and part of the last exon. Asterisks indicate editing sites. (E) Quantification of the level of LEM-2 protein in WT and ADAR mutant worms after *lem-2* RNAi relative to empty vector. LEM-2 protein expression levels were normalized to actin protein levels. The experiment was repeated at least 3 times with similar results. Error bars represent SD. * P-value < 0.05 calculated by unpaired student’s t-test.

### Editing significantly reduces the efficiency of exogenous RNAi

To further investigate the antagonistic effect of RNA editing on RNAi *in vivo*, we searched for an RNA-edited gene that exhibits a pronounced phenotype. We chose *egl-2* as a candidate. *egl-2* undergoes RNA editing at multiple sites at its 3’UTR (Goldstein et al., 2017), is highly enriched in secondary siRNAs in ADAR mutants (Figure 1D, Supp Figure 1D), and *egl-2* mutant strains have very noticeable phenotypes such as egg laying defects leading to bloated worms ((Loria et al., 2003; Trent et al., 1983; Troemel et al., 1999)). Since downregulation of *egl-2* by RNAi does not exhibit any phenotype in wildtype animals (Kamath et al., 2003; Shephard et al., 2011; Sönnichsen et al., 2005), our hypothesis was that RNA editing prevents primary siRNA generation in wildtype worms, causing the ineffectiveness of RNAi. If our hypothesis was correct, we expected to observe significant downregulation of *egl-2* in the ADAR mutant background when performing RNAi, as well as severe phenotypes in comparison to the wildtype. To test this prediction, we performed an RNAi experiment on wildtype and ADAR mutant worms using a bacterial feeding approach with a trigger specific to the RNA editing-rich region or to the entire 3’UTR of *egl-2* (Figure 4A). We found that indeed, RNAi on wildtype worms did not yield severe phenotypes, while RNAi on the ADAR mutant strains resulted in similar severe phenotypes detected in *egl-2* mutant strains (Figure 4B, Supp Figure 5). We confirmed the downregulation of *egl-2* RNA in ADAR mutant worms and non-significant downregulation in wildtype worms by real-time PCR (Figure 4C). A small fraction of the *egl-2* phenotype was also observed when we performed RNAi with the background plasmid that we used for cloning, which does not produce dsRNAs against a *C. elegans* gene (e.g., empty vector) on ADAR mutant worms (Figure 4B). This background phenotype was expected because we found enrichment of secondary siRNAs in ADAR mutant worms (Figure 1D, Supp Figure 1D) in the absence of exogenous triggered RNAi. RNAi against a region in *egl-2* RNA that does not undergo editing did not yield a significant *egl-2* phenotype in ADAR mutant worms compared to empty vector (Supp Figure 6). In addition, the enrichment in RNAi effectiveness was not because of hypersensitivity to RNAi in ADAR mutants, since this enrichment was not seen in the RNAi hypersensitive mutant, *rrf-3 (pk1426)* (Figure 4B).

**Figure 5.**
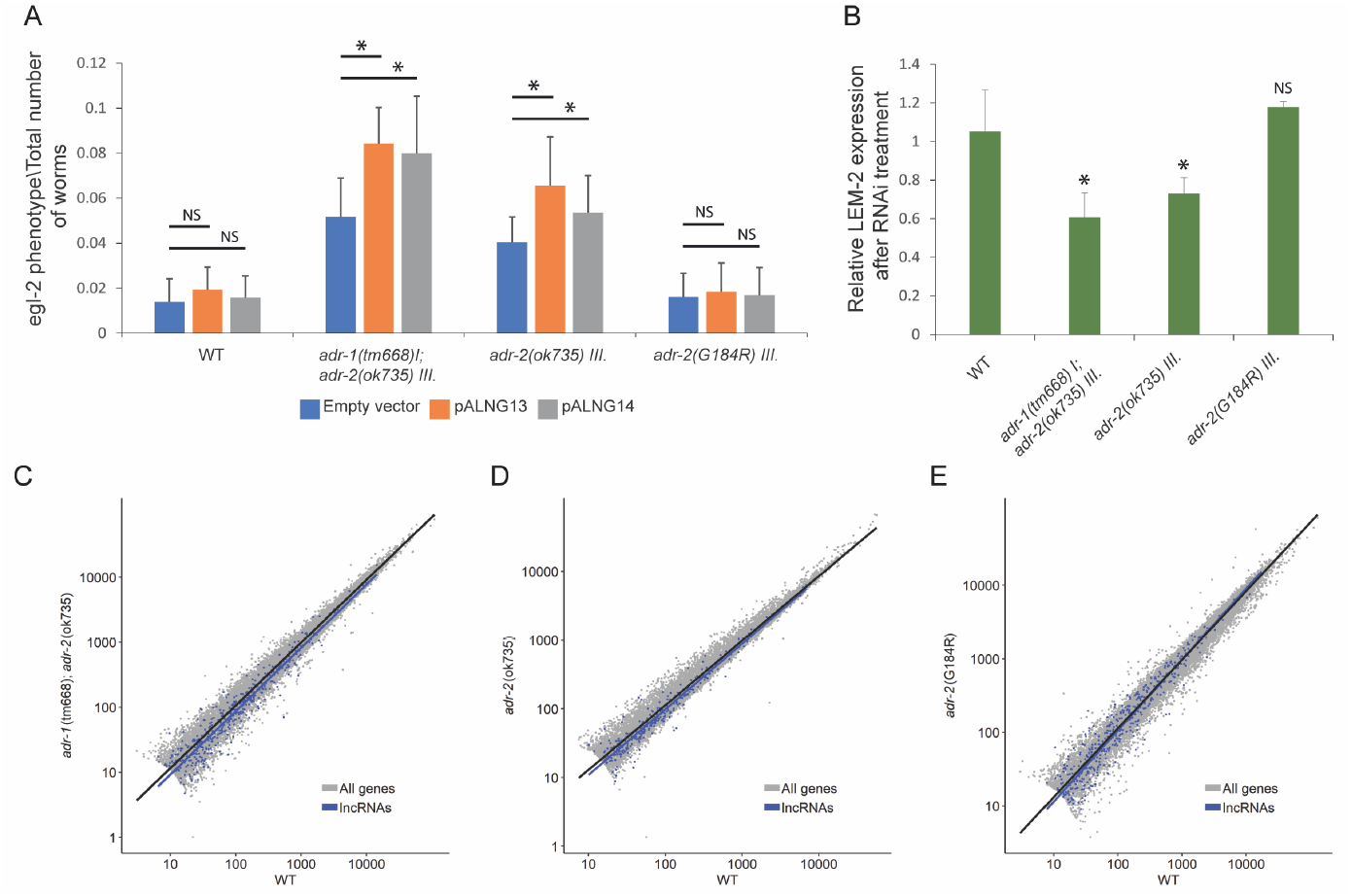
Mutation that eliminates the deamination ability of ADR-2 does not prevent RNAi. (A) Quantification of the fraction of *egl-2* phenotype from total worms observed in the RNAi experiments. Abnormal phenotypes that were scored include bloated worms, exploded worms, egg retention, and bag of worm phenotypes. *adr-2* (G184R) is *adr-2* mutated in its deamination domain; P-values were calculated from at least 3 biological replicas, NS for non-significant, *P-value < 0.05. (B) Quantification of the level of LEM-2 protein in wildtype (WT) and ADAR mutant worms. Protein levels were measured following RNAi treatment against *lem-2* and compared to those in worms treated with an empty vector. All values were normalized to Actin levels. The experiment was repeated at least 3 times with similar results and SD is presented. Statistical analysis was done by comparing each mutant strain with the wildtype. * P-value < 0.05 calculated by unpaired student’s t-test. (C-E) Log scale plot presenting normalized counts of genes and lncRNA from RNA-seq data at embryo stage in wild-type (WT) worms and ADAR mutant (*adr-1* (tm668); *adr-2* (ok735)), *adr-2* (ok735), *adr-2* (G184R)) worms. Every dot represents a gene. Gray dots indicate all genes. Blue dots indicate lncRNA. The black line is the regression line for all genes. The blue line is the regression line for lncRNA.

**Figure 6.**
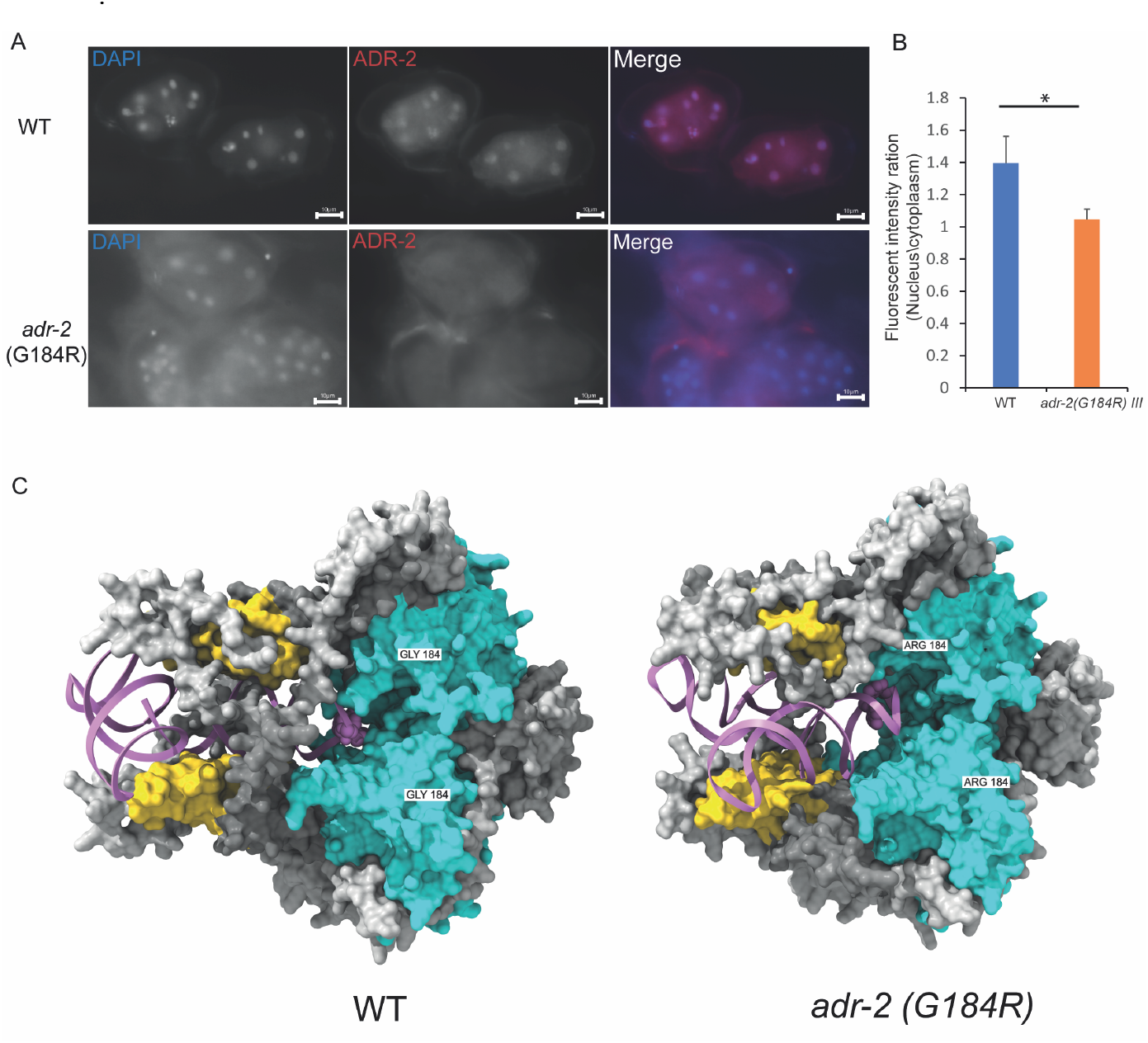
Mutation in the deamination domain of ADR-2 enhances ADR-2 cytoplasmic localization. (**A**) Immunostaining of wildtype and *adr-2*(G184R) embryos using ADR-2 antibody. Anti-ADR-2 antibody staining is shown in red in the merge images. Nuclei were visualized with DAPI staining and are shown in blue in the merge images. Scale bar, 10 µm. (B) The fluorescence intensities of ADR-2 in the nucleus and in the cytoplasm were quantified. The graph shows the mean nuclear/cytoplasm ratios in at least 3 cells in each embryo (at least 3) in 3 biological replicates. *P < 0.05. (C) AlphaFold3 ADR-2 X2 (silver) ;ADBP-1X2 (white) tetrameric prediction based on (Mu et al., 2025) with *egl-2* RNA (pink). On the left is ADR-2 wildtype, and on the right ADR-2 with G184R mutation. Pink spheres in the RNA are editing sites. In yellow is ADR-2 RNA binding domain, and in blue ADR-2 deaminase domain. ADR-2 184 amino acid is marked in white.

To further confirm that loss of RNA editing can restore exogenous RNAi efficiency, we explored whether loss of RNA editing can also affect the protein levels through its antagonistic relation to RNAi. We, therefore, triggered RNAi on a gene that is heavily edited at its 3’UTR, *lem-2*, which is also enriched in primary siRNAs in ADAR mutant worms (Figure 1C, Supp Figure 1C) and monitored effects on protein levels by immunoblotting lysates prepared from wildtype and ADAR mutant worms (Figure 4D,E, Supp Figure 7). Consistent with our other observations, we also observed a significant downregulation of LEM-2 protein in ADAR mutant worms compared to wildtype worms (Figure 4D,E, Supp Figure 7). We concluded that RNA editing probably protects endogenous genes from externally triggered RNAi or viral-triggered RNAi.

### ADAR binding without deamination activity prevents RNAi

After establishing that ADARs prevent exogenous RNAi, we wanted to explore how this prevention occurs. One possibility was that inosines prevent Dicer from binding either by changing the conformation of the dsRNA and reducing the binding efficiency or because Dicer cannot bind inosines efficiently. Another possibility was that by directly binding the dsRNA, ADARs preclude Dicer from binding (reviewed in (Ganem and Lamm, 2017)). The latter option seemed less likely as ADARs were shown to localize mainly in the nucleus, while RNAi occurs mainly in the cytoplasm, although the enzymes from both processes were found to be in both compartments (Desterro et al., 2003; Drake et al., 2014; Eliad et al., 2024; Fritz et al., 2009; Hundley et al., 2008; Ohta et al., 2008). To determine whether the adenosine-to-inosine change prevents RNAi, we used a strain that harbors a point mutation in the deamination domain of ADR-2, resulting in the inability to deaminate RNA while leaving the dsRNA binding domain of ADR-2 intact, *adr-2 (G184R)* (Deffit et al., 2017). As expected (Deffit et al., 2017), *egl-2, lem-2* and other edited genes do not undergo editing in this strain (Supp Figure 8). Surprisingly, we found that triggering RNAi against *egl-2* in this mutant strain does not result in a pronounced *egl-2* phenotype by silencing (Figure 5A), while a deletion in *adr-2, adr-2 (ok735)*, which does not generate a protein, shows enrichment in the phenotype (Figure 5A). To further confirm that even without editing ability, ADR-2 protects dsRNAs from RNAi, we tested the protein levels of *lem-2* RNAi after triggering RNAi in this mutant. Similar to the *egl-2* RNAi phenotype, we did not observe a change in LEM-2 protein levels in the *adr-2* deamination mutant (*adr-2(G184R)*, Figure 5B), while a significant reduction in the protein levels was observed in *adr-2* deletion mutant (*adr-2(ok735)*, Figure 5B).

We previously showed a significant reduction in the level of lncRNAs in ADAR mutant compared to wildtype ((Goldstein et al., 2017)) by RNA sequencing. We speculated that this reduction is a result of the enhancement of the RNAi process by removing the inhibition of RNA editing in ADAR mutant, and indeed, we observed an increase in the number of siRNAs aligned to lncRNAs in ADAR mutant (Goldstein et al., 2017). To test if the lncRNAs expression is attenuated in *adr-2* deamination mutant, we generated RNA-seq libraries or used our published RNA-seq libraries from wildtype, *adr-1*;*adr-2* double mutant, *adr-2* deletion, and *adr-2* deamination mutant worms at the embryonic developmental stage (Ganem et al., 2019; Goldstein et al., 2017). As expected, lncRNA expression is significantly reduced in *adr-1*;*adr-2* mutant worms compared to wildtype worms (Figure 5C, P-value < 3e-16 calculated by Welch two-sample t-test). We also observed a significant reduction in lncRNA expression in the *adr-2* deletion (Figure 5D, P-value <7e-10), while no reduction was observed in *adr-2* deamination mutation (Figure 5E, P-value non-significant, Supp Table 1).

These results suggest that ADR-2 binding to RNA, even without deamination activity, prevents processing by RNAi. In addition, our results indicate that the first step of the RNA processing by RNAi, e.g. generation of primary siRNAs, is the only step that is inhibited. This step mainly occurs in the cytoplasm by Dicer and other RNAi components, while ADR-2 mainly acts in the nucleoplasm. Although ADR-2 mainly resides in the nucleoplasm it was shown to shuttle to the cytoplasm and its localization is regulated by ADBP-1 protein (Eliad et al., 2024; Ohta et al., 2008). Therefore, it is possible that the deamination mutation in ADR-2 protein changes its localization and by that inhibits RNAi directly in the cytoplasm. To better understand ADR-2 localization and activity, we used an antibody against ADR-2. We immunostained ADR-2 and stained DNA using DAPI in wildtype and *adr-2* deamination mutant embryos, similarly to (Eliad et al., 2024) (Figure 6A). In wildtype embryos, most ADR-2 protein was localized to the nucleoplasm, while in the *adr-2* deamination mutant, there was significant localization of the protein in the cytoplasm (Figure 6B). The difference in the localization of ADR-2 protein in the mutant was not because of a change in the protein levels, which was shown to be the same in wildtype and *adr-2* deamination mutant (Deffit et al., 2017). We also did not observe a change in the binding ability of ADR-2 to the RNA in the wildtype and *adr-2* deamination mutant (Supp Figure 9). These results suggest that ADR-2 binding to the RNA, even without editing, prevents RNAi from processing the dsRNA, probably by not leaving enough space for RNAi components to bind the dsRNA.

Recently, the structure of ADBP-1 and ADR-2 tetrameric protein complex was resolved by cryo-EM (Mu et al., 2025). They showed, as was estimated by AlphaFold2 (Eliad et al., 2024), that ADBP-1 binds ADR-2 near its deamination domain. ADR-2 can edit RNA independently of ADBP-1, when it resides in the cytoplasm (Eliad et al., 2024; Mu et al., 2025). In addition, (Mu et al., 2025) found that ADBP-1 increases ADR-2 RNA binding. They also suggested that G184R mutation affects the RNA binding of ADR-2. To further estimate the effect of G184R mutation, we used AlphaFold3 to model the change in structure using ((Mu et al., 2025); PDB:8ZEP) ADR-2 and ADBP-1 tetrameric structure with 80 RNA nucleotides from *egl-2* 3’UTR that undergo editing (Figure 6C). Our model suggests that the mutation increases the positive electrostatic field (Supp Figure 10A), causing a stronger RNA binding and stability of the complex with the RNA as was calculated by pyDock (Supp Table 2). On the other hand, the mutation causes two regions in the deaminase domain to be less organized and probably more flexible with lower pLDDT values (Supp Figure 10B-D). We hypothesize that this flexibility likely prevents active deamination while facilitating stronger binding to the RNA.

Therefore, the results suggest that in wildtype worms, ADR-2 binds and then deaminates the RNA in the nucleus. After deamination, in most cases, ADR-2 is released from the RNA, and the deamination itself protects it from RNA interference (RNAi). However, in the ADR-2 deamination mutant, ADR-2 resides mainly in the cytoplasm. ADR-2 binds the RNA but cannot deaminate it. Therefore, it is not released from the RNA, preventing Dicer from binding to the dsRNA, and protecting it from the RNAi machinery (see graphical abstract).

## Discussion

The important function of A-to-I RNA editing in regulating the immune system, specifically the interferon reaction, in mammals (Reviewed in (Quin et al., 2021)) is well established. The main paradigm is that ADARs edit endogenous dsRNA in the nucleus to select them from viral/exogenous dsRNA to prevent mis-activation of the interferon reaction against self-produced RNA. ADARp150, ADAR1 cytoplasmic isoform, also regulates the interferon reaction in the opposite direction. It prevents overactivation of the interferon reaction (Quin et al., 2021). *C. elegans* does not have the interferon mechanism but rather has a very extensive RNAi mechanism. There are several functional legs for RNAi. One of them is defense against exogenous RNA by detecting dsRNA and generating siRNAs that lead to silencing or degradation (Grishok, 2013). Several studies showed an antagonistic relationship between A-to-I RNA editing and RNAi (Goldstein et al., 2017; Reich et al., 2018; Warf et al., 2012; Wu et al., 2011), suggesting that A-to-I RNA editing in *C. elegans* regulates the immune system similarly to mammals. e.g. *C. elegans* ADARs edit endogenous dsRNAs in the nucleus to prevent Dicer and the RNAi machinery from silencing them, while exogenous/viral RNAs are not edited and are targeted for silencing by RNAi. Evidence comes from our previous study that shows that in the absence of ADRs, downregulation of edited genes is observed (Goldstein et al., 2017) with a parallel enhancement in the number of siRNAs against edited genes (Goldstein et al., 2017; Wu et al., 2011). In addition, ADARs were also shown to silence retrotransposons in *C. elegans* (Fischer and Ruvkun, 2020), and that the downregulation of dsRNAs in the absence of ADARs is dependent on several components of RNAi (Reich et al., 2018).

RNAi processing has two main steps. In the first step, the dsRNA is recognized, and primary siRNAs are generated. This step is mainly catalyzed by DICER protein. In the second step, primary siRNAs are used to amplify the reaction by generating secondary siRNAs. This step is mainly catalyzed by RdRP proteins (Grishok, 2013). Here we sought to define which of these steps, or both, are inhibited by RNA editing by generating and analyzing small RNA-seq data enriched for either primary or secondary siRNAs. As we showed before (Goldstein et al., 2017; Wu et al., 2011), we found that in the absence of RNA editing, the number of siRNAs against edited genes, both primary and secondary, is significantly enhanced but not against other genes. This is in line with the notion that RNA editing inhibits RNAi, and when this inhibition is removed, RNAi processes the dsRNA and leads to downregulation of genes. Examining the sizes of the siRNAs aligned to editing sites, we found that both primary and secondary siRNAs were enriched for 23-mer siRNAs in ADAR mutants compared to wildtype but other sizes were not affected. This suggests that the main RNAi pathway that is inhibited by ADARs is viral RNAi, which produces 23-mer siRNAs (Ashe et al., 2013), but it is also possible that endogenous 23-mers might be affected as well (Knittel et al., 2024). We found that the generation of primary siRNAs is significantly prevented at RNA editing sites by the editing process, while the generation of secondary siRNAs by RdRPs can still occur (Figure 2F). Furthermore, the finding that secondary siRNAs contain editing changes allowed us to deduce that only the first step of recognition of the dsRNA by Dicer and other components is inhibited by RNA editing and that RdRPs can still process edited transcripts to generate secondary siRNAs. If primary siRNAs are generated from a nearby location, secondary siRNAs can be generated and will have the editing change; therefore, they might trigger gene silencing at a different location that is identical to the modified siRNAs, but they are not enough to silence the edited genes. In addition, the fact that secondary siRNAs contain editing changes allowed us to deduce the sequence of events that happen when dsRNA is triggered for processing. In other words, it allowed us to figure out whether siRNAs are generated before editing takes place or after it. If siRNAs were generated before editing takes place, they could not contain the editing change. Our results are consistent with the latter option, namely that siRNAs are generated after editing events and therefore can bear the editing changes.

It is not clear why specific genes are edited and others are not. One possibility is that genes and dsRNAs highly similar to viral RNA can be silenced by trans-silencing when these viruses enter the cell and are targeted by RNAi. To test this hypothesis, we generated a system that shows *in vivo* that when triggering exogenous RNAi against editing sites, specifically the 3’UTR of *egl-2* or *lem-2*, silencing does not occur. But when removing editing, by using ADAR mutants, we observe that the downregulation effect leads to RNA downregulation, less protein production, and phenotypes that are similar to when the gene is absent (Figures 4,5). Another support for this hypothesis comes from Reich et al., that showed that many genes that are differently expressed during Orsay virus infection are also differentially expressed in the absence of ADARs and the RdRP, *rrf-3* (Reich et al., 2018). We could not find similarity of *egl-2* and *lem-2* RNA to known *C. elegans* viruses, however not many are known.

To study how editing inhibits RNAi, we used an *adr-2* mutant that is catalytically inactive and cannot edit because of a point mutation, however it can still bind RNA (Deffit et al., 2017). We tested its ability to inhibit RNAi in an exogenous triggering reaction. Because ADR-2 mainly resides in the nucleus, by our immunostaining results, and because of indications *in vitro* that Dicer is inhibited by inosines directly (Scadden and Smith, 2001), we expected that triggering exogenous RNAi in this mutant against edited sites would lead to activation of RNAi and downregulation. Our results were surprising, showing inhibition of RNAi in editing sites similarly to wildtype worms in all systems tested. Additionally, we did not observe downregulation of lncRNAs in this mutant as in *adr-2* deletion strain, which also suggests that inhibition of RNAi occurs in this strain. It is possible that there are still residues of editing in *adr-2* deamination mutant, however we could not observe them in the regions that we triggered or in other highly edited genes. It is more probable that the ability of ADR-2 to still bind the dsRNA in the deamination mutant causes RNAi inhibition.

This possibility is confusing because deamination mainly occurs in the nucleoplasm and silencing by RNAi mainly occurs in the cytoplasm. However, upon analysis of the localization of ADR-2 by immunostaining ADR-2 in *adr-2* deamination mutant, we found that in contrast to wildtype worms, the ADR-2 G184R deamination mutant protein highly resides in the cytoplasm. Therefore, it is possible that ADR-2, which is in the cytoplasm, protects the dsRNA from RNAi. It was shown that ADAR1 in mammals binds Dicer through its second dsRNA binding domain (Ota et al., 2013). It is possible that a similar interaction occurs in *C. elegans* in wildtype worms although it is not clear in which cellular compartment. ADBP-1 protein was shown to regulate ADR-2 cellular localization, and therefore, it is possible that ADBP-1 is involved in the regulation of RNAi as well (Eliad et al., 2024; Ohta et al., 2008). Indeed, transgene silencing was observed in *adbp-1* mutant worms (Ohta et al., 2008). To test this further, we used the known tetrameric structure of ADR-2 and ADBP-1 (Mu et al., 2025) to model the changes the G184R mutation causes to the structure. The model does not support a change in ADR-2 and ADBP-1 binding, however the mutation seems to cause a stronger binding to the RNA with a more stable structure. RIP experiments also showed binding to the RNA when ADR-2 is mutated. Additionally, the mutation results in a more flexible structure at the deamination sites, which likely causes the enzyme’s inability to deaminate and a dominant negative effect by preventing the release of RNA. We suggest that the sequence of events leading to editing a specific site is that ADR-1 leads ADR-2 to the deamination site, next ADR-2 binds the dsRNA through its RNA binding domain, deaminate the dsRNA, and then releases from the dsRNA, free to deaminate another target. When a mutation occurs in the deamination domain of ADR-2, rendering it unable to deaminate after binding to RNA, ADR-2 cannot deaminate the RNA and fails to release from it. Then, the ADR-2-RNA complex in the cytoplasm interferes with DICER binding to the dsRNA and generating primary siRNAs. This does not exclude the possibility that inosines on the dsRNA also inhibit DICER binding in wildtype worms. While in worms with *adr-2* full deletion, both deamination and inhibition by binding do not occur, with no protection against RNAi, and downregulation of edited genes is observed. We previously showed that ADR-2 is functional in the cytoplasm and can edit RNA (Eliad et al., 2024). One possibility is that mutated ADR-2, with or without ADBP-1, before entering the nucleus, binds dsRNA in the cytoplasm and cannot release it, preventing ADR-2 from entering the nucleus. Another possibility is that after entering the nucleus with ADBP-1, ADR-2 binds RNA, cannot release the RNA, and shuttles with it to the cytoplasm.

To conclude, we found that the generation of primary siRNAs is significantly prevented in RNA editing sites by the editing process, while the generation of secondary siRNAs by RdRPs can still occur. This interference with the RNAi process also affects the efficiency of exogenous RNAi on edited genes, probably to protect them from being recognized by RNAi as viral RNA and possibly to protect them from trans-silencing. We also showed that ADAR binding is needed for the prevention of exogenous RNAi by ADARs, suggesting that ADR-2 dsRNA binding might have a different function from its deamination function.

## Materials and Methods

### Maintenance and handling of *C. elegans* strains

The following strains were used in this study: Bristol N2, BB4 *adr-1 (gv6)* I; *adr-2 (gv42)* III) (Tonkin et al., 2002), BB19 *adr-1 (tm668)* I, RB886 *adr-2 (ok735)* III, BB21 *adr-1 (tm668)* I; *adr-2 (ok735)* III (Hundley et al., 2008); HAH4 *adr-2 (gk777511)* (Deffit et al., 2017). All strains were grown at 20°C as described in (Brenner, 1974). Worms were synchronized to the embryo stage by using sodium hypochlorite. For RNA extraction, embryos were frozen into pellets with liquid nitrogen.

### Generation of siRNAs libraries and processing

Embryos frozen pellets were ground into powder with a liquid nitrogen chilled mortar and pestle. RNA in high and low molecular weight fractions was extracted by mirVana kit (Ambion). Small RNA sequencing libraries were prepared from the low molecular weight fraction using QsRNA-seq method (Fishman et al., 2018). Secondary siRNAs were sequenced using the same method by adding an initial step, incubation with 5U of RppH enzyme (NEB) for 30 min at 37^0^C to remove two 5’-phosphates (Fishman et al., 2018). The sequences generated have 4nt-barcode and 8N for UMI. RNA sequences were trimmed to remove adapter sequences as described in (Fishman et al., 2018). In short, sequences were first de-multiplexed according to the 4nt barcode. Next, the 3’adapter sequences were trimmed off by scanning from the 3’-end of the sequence for the first instance of the adapter sequence by increments of 1nt. Then, identical sequences were merged and barcode and UMI were removed. Primary siRNA and secondary siRNA libraries were filtered to include only sequences of sizes 19-27bp. The sequences were then aligned to the WS220 (Wormbase, www.wormbase.org) genome or gene datasets using Bowtie (Langmead et al., 2009). The amount and size of siRNAs aligned to editing sites were then estimated. Fold change and p-adj value (P-value after BH correction) were estimated using DEseq2 (Anders and Huber, 2010) package in R (http://www.r-project.org).

### Detailed bioinformatics analysis

#### Site resolution analysis

Fastq files were aligned to the entire *C. elegans* chromosomes (WS220) using bowtie (Langmead et al., 2009), allowing up to two mismatches. This version was chosen to match the sites in (Ganem et al., 2019). The aligned sam files were converted to pileups using samtools (Li et al., 2009). Each siRNA pileup file was split into two files, sense and antisense based on the gene orientation. We assume that the anti-sense file is comprised of siRNAs, while the sense file is comprised mostly of degradation products. We then counted the number of reads per nucleotide in the antisense files and analyzed the results through DESeq2.

#### Editing percent analysis

All biological repeats of every siRNA antisense pileup were merged. The resulting pileups were filtered according to the non-repetitive editing sites and 3’UTR edited sites from (Ganem et al., 2019; Goldstein et al., 2017). The editing percentage was calculated according to the pileup base string.

#### Priming distance analysis

Secondary siRNA pileup files and primary siRNA sam files were merged according to sample type. For each secondary siRNA nucleotide that was present in a pileup, we looked for the closest primary siRNA read that appeared before the secondary siRNA nucleotide and calculated the distance between them. P-values for the distance between primary and secondary siRNAs were computed using the anderson_ksamp function of the scipy (Jones et al., 2001) statistical computation package.

### mRNA-seq library preparation and processing

Embryo frozen pellets were ground into powder with liquid nitrogen chilled mortar and pestle. RNA was extracted by using Direct-zol RNA MiniPrep Plus (ZYMO). The RNA was treated with TURBO DNA-free Kit (Invitrogen by Thermo Fisher Scientific, AM1907). mRNA sequencing libraries were prepared by using TruSeq RNA kit from Illumina and sequenced by Illumina HiSeq 2500. For analysis, libraries were generated here or were taken from our published work (GSE83133, GSE110701). The reads were aligned to *C. elegans* WS220 transcriptome using Bowtie (Langmead et al., 2009) . Fold change and p-adj value (P-value after BH correction) were estimated using DEseq2 (Anders and Huber, 2010) package in R (http://www.r-project.org). To compare the expression of lncRNA and 3’UTR edited genes to all the expressed genes, we used the Welch two-sample t-test between chosen two groups (two-sided).

### RNAi and phenotypic assay

RNAi feeding constructs were generated by cloning *egl-2* and *lem-2* 3’UTR into L4440 (Fire lab vector kit) feeding vector using XmaI enzyme (NEB®). Plasmids PALNG013 contain the entire *egl-2* 3’ UTR using primers AL_NG25 and AL_NG26. PALNG014 contains partial *egl-2* 3’ UTR using primers AL_NG25 and AL_NG27. Plasmid PALNG020 contains the entire lem-2 3’ UTR, including 34bp upstream of the 3’UTR using primers AL_NG54 and AL_NG55 (the list of primers is in the supplementary materials). The plasmids were transformed into E. coli HT115 bacteria and were cultured overnight at 37°C in LB media containing 100 μg/ml ampicillin. The cultured bacteria were seeded onto *C. elegans* growth media plates (NGM) containing 100μg/ml ampicillin and 10 μg/ml IPTG and incubated overnight at RT. Synchronized embryos were placed on the plates and incubated for 72-96 hours at 20°C before scoring or collected for real-time PCR or western blot. For phenotypic characterization, worms were then scored using a Nikon SMZ745 zoom stereomicroscope for *egl-2* phenotypes, including Pvl, bag of worms, bloated worms, expulsion defective, fat, and defecation variant. MT1444 *egl-2(n693)*V strain was used as a reference. Worms presenting the phenotypes were counted relative to the total number of worms in each plate. P-value was calculated by two tailed t-test from 3 biological replicas.

### Western blot assay

Worms after *lem-2* RNAi treatment were collected and then washed five times using M9 buffer until the sample was clear. Sample loading buffer (SLB) was used to lyse the worms at 100°C for five minutes. Protein extracts, separated by SDS-PAGE and transferred onto cellulose membrane, and then immunoblotted with antibodies against LEM-2 (Rabbit anti-LEM-2, 1:2000, 48540002 Novus biologicals) or against actin (Mouse anti-actin monoclonal, clone, 1:400, 0869100 MP BIOMEDICALS). LEM-2 protein was detected with goat anti-rabbit IgG antibody (Peroxidase AffiniPure Goat Anti-Rabbit IgG (H+L), 1: 15000, 111035003 Jackson immunoresearch), Actin protein was detected with goat anti-mouse IgG antibody (Peroxidase AffiniPure Goat Anti-Mouse IgG (H+L), 1: 15000, 115035003 Jackson immunoresearch), and visualized with the EZ-ECL Western blotting substrate (EZ-ECL Chemiluminescence Detection Kit for HRP, 20-500-120 Biological Industries), according to the provided protocol. Proteins chemiluminescence signal was detected and quantified using ImageJ image analysis program. Actin protein was used as a control for normalization.

### Real-time PCR

RNA (9ug) was treated with DNase using TURBO DNA-free kit (Invitrogen) and then 500ng was reverse transcribed by Q-SCRIPT flex cDNA synthesis kit (Quanta Biosciences) using random primers, according to the manufacturer’s protocol. qPCR primers were designed using Primer designing tool (http://www.ncbi.nlm.nih.gov/tools/primer-blast). When possible, primers spanning exon junctions were chosen. qPCR reactions were prepared in triplicate using PerfeCTa SYBR Green FastMix ROX (Quanta Biosciences) according to the manufacturer’s protocol and run on CFX-96 Real Time system (Bio-Rad). The results were analyzed using Bio-Rad CFX manager software. Gene expression was calculated by normalizing to reference genes (ΔΔCq). The melting curve confirmed the existence of a single product in each reaction. All NTC controls were negative. All NRT controls were either negative or not significant (<3% of total product). All the primers used for the RT-PCR reactions are listed in the supplementary materials, including the calibrator genes.

### Immunostaining, Microscopy, and intensity quantification

Adult *C. elegans* worms were fixed and prepared for immunostaining according to (Margalit et al., 2005). Primary Rabbit anti-ADR-2 (Rajendren et al., 2018) was used at 1:50 dilution for indirect immunostaining. Secondary Donkey anti-Rabbit IgG (H+L) Highly Cross-Adsorbed Secondary Antibody, Alexa Fluor® 568 (Life Technologies) was used at 1:200 dilutions. 4′, 6-diamidino-2-phenylindole (DAPI) were used at 1:1000 for DNA staining. Images were taken using a fluorescent microscope (Nikon Eclipse Ni-E) with CCD camera (ANDOR iXon Ultra) with a 100x oil immersion objective lens. The fluorescence images were analyzed with the ImageJ image analysis program. By using ROI measurement tool, the fluorescence intensities of the cytoplasmic regions and the nucleus were measured and averaged. The nuclear-cytoplasmic ratio of ADR-2 localization in each strain was determined as the average measurement of 24 cells from eight different embryos. The statistical analysis was performed using two-sample, unequal-variance, heteroscedastic T-test.

### ADR-2 mutation complex modeling

We modeled the WT ADR-2 and ADR-2 G184R mutant in complex with ADBP-1 and RNA with AlphaFold3 (Abramson et al., 2024). The tetrameric structure was taken from a recent paper (Mu et al., 2025) (PDB: 8ZEP). The RNA we used for the model was from *egl-2* 3’UTR that we used for the RNAi ( see supplementary materials for the sequence). To compute the interface binding energy between the RNA molecule and all 4 peptides in the structure, we used pyDock (Cheng et al., 2007).

## Acknowledgments

We want to thank Tal Elfand for helping with the bioinformatics analysis. This work was funded by The Israel Science Foundation (grant 1080/23 to ATL), the Binational Israel-USA Science Foundation (grant No. 2023134 to ATL and HAH), the NSF-BSF Molecular and Cellular Biosciences (MCB) (grant No. 2018738 to ATL and HAH), and the National Institutes of Health (R35GM156459). Some strains were provided by the CGC, which is funded by NIH Office of Research Infrastructure Programs (P40 OD010440).

## declaration of interests

The authors declare no competing interests.

## Notes

### Competing Interest Statement

The authors have declared no competing interest.

### Summary of Updates

Added a graphical abstract to explain the mechanism.

