## Supplemental figures and methods for "RNA binding by ADARs prevents RNA interference from attacking self-produced dsRNA"

**Supplementary Figures and Methods**

**Supplementary Material and Methods**

**Primers used in this study**

**Primers used for sequencing**

*egl-2* forward - ATTTGTGATATGTAAACTTCATTG

*egl-2* reverse - CAGTTTTAATAAATAAAAACGGC

*lem-2* forward - GTGGATCGGAAATCAGTCTC

*lem-2* reverse - CGAGCAAAAATCCATCG

*F48E8.4* forward - CTTCTAGTCCCGCCAAATTTATG

*F48E8.4* reverse - CAGTTGAAGTTATTCCACGACCC

*C54D1.5* forward - TGACGAGGACGTAAAGGTGGC

*C54D1.5* reverse – CGTATTACTACTCAGATGACC

**Primers used for real-time PCR**

AL_NG_60 *egl-2* exon 6 Forward CGCAAAGTCAAAGCAGCAGT

AL_NG_61 *egl-2* exon 6 Reverse ACCTGATTGAAGTTACTGGATGTG

AL_NG_62 *egl-2* exon 6 Forward TAGCACGGATCGCAAAGTCA

Primers used for calibration:

AL-AF-28 *orai-1* Forward TGCAGCTGGAATAGATCCCC

AL-AF-29 *orai-1* Reverse TTGGAGCATTTTCCACGGGT

AL-AF-30 *fce-1* Forward TCTGGGCCATGAATTGGGTC

AL-AF-31 *fce-1* Reverse GCCGAATCCTTGATAGAGGGC

AL-AF-32 *scpl-4* Forward CGCCGGAAACGAATGGAAAG

AL-AF-33 *scpl-4* Reverse ACGGTGCCAAGAAAGAACCA

**Primers used for cloning**

AL_NG_54 *lem-2* forward gtcacccgggGTGGATCGGAAATCAGTCTC

AL_NG_55 *lem-2* reverse gtcacccgggCGAGCAAAAATCCATCG

AL_NG_25 *egl-2* forward gatcCCCGGGcacatggatccttttccaa

AL_NG_26 *egl-2* reverse gatcCCCGGGagcatctgccaaaatatttattta

AL_NG_27 *egl-2* forward gatcCCCGGGggcactgcaactttttcctc

**RNA used for the structural model**

5'-
cuuuuuccucacgagggacuaggaaaagugguuucuaggccauggcugaggggccgacaaguuucagcggucauuuauc
u -3'

**DNA and RNA Sanger sequencing**

To obtain cDNA, extracted RNA (DirectZol , Zymo) was treated with DNase I (Ambion), and then a reverse transcriptase reaction was performed with Maxima First Strand cDNA Synthesis kit (Thermo Scientific), using oligo (dT)18 and random hexamer primers. DNA was extracted using Phire Tissue Direct PCR Master Mix (Thermo Scientific). The amplification products were directly sequenced by Sanger sequencing.

**RNA immunoprecipitation (RIP) assay**

RNA immunoprecipitation assays for ADR-2 were performed as previously described in (Deffit et al., 2017; Rajendren et al., 2018). Briefly, after washing with IP buffer (50 mM HEPES [pH 7.4]; 70 mM K-Acetate, 5 mM Mg-Acetate, 0.05% NP-40, and 10% glycerol), worms were subjected to 3 J/cm2 of UV radiation using the Spectrolinker (Spectronics, Westbury, NY) and stored at −80°C. To obtain cell lysates, frozen worms were ground with a mortar and pestle on dry ice. After thawing, the lysate was centrifuged, and protein concentration was measured with Bradford reagent (Sigma-Aldrich). Five milligrams of extract were added to ADR-2 antibody (Cocalico Biologicals- IU529) coated anti-Rabbit IgG magnetic Dynabeads (Fisher). After incubation for 1 hr at 4°C, the beads were washed with wash buffer (WB: 0.5 M NaCl, 160 mM Tris-HCl [pH 7.5]) resuspended in low-salt WB (0.11 M NaCl), 1 µl RNasin (Promega, Madison, WI), and 0.5 µl of 20 mg/ml proteinase K (Sigma-Aldrich) and incubated at 42°C for 15 min to degrade protein and release bound RNA. Protein samples were subjected to SDS-PAGE and western blotting with ADR-2 antibody (Cocalico Biologicals-IU529). Gene-specific primers were used to synthesize complementary DNA (cDNA), and quantitative real-time PCR with gene-specific primers was performed to quantify mRNA abundance. Statistical significance between the samples with three biological replicates was calculated with student t-test.

### Supplementary Figures

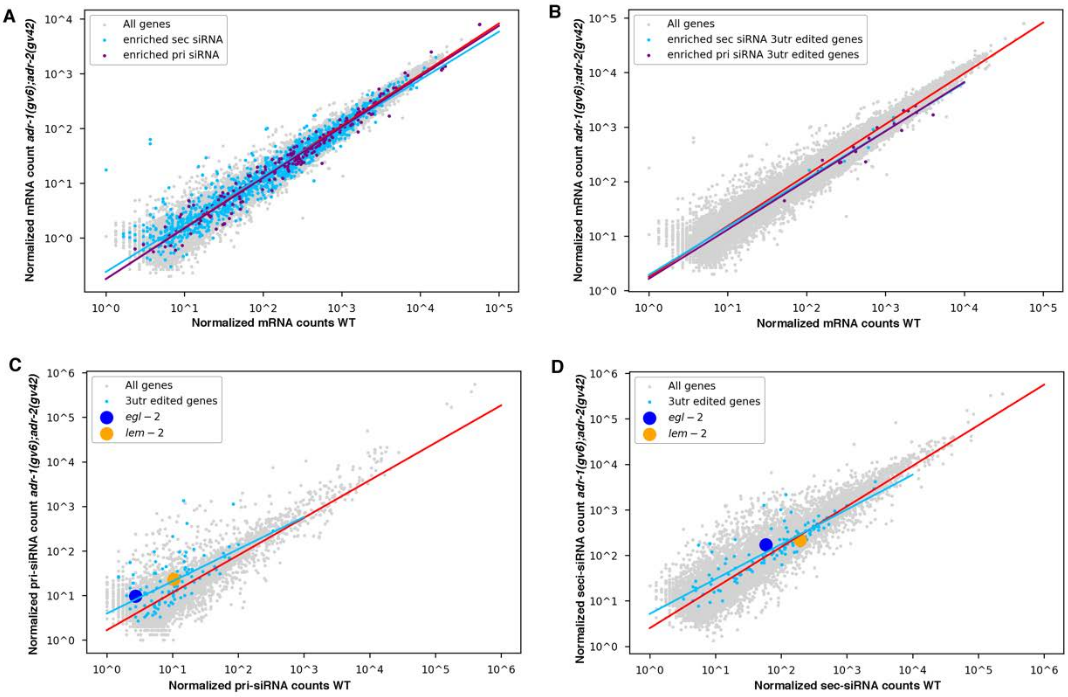


**Supplementary Figure 1. Genes edited at their 3'UTR are siRNAs enriched in ADAR mutant worms**. Log scale plots presenting normalized mRNA sequence counts (A,B), primary siRNAs sequence counts (C), or secondary siRNAs sequence counts (D) of genes from at least 3 biological replicas in wildtype (N2) worms compared to ADAR mutant worms (*adr-1 (gv6) ; adr-2 (gv42))*. Every dot in the graphs represents a gene. The red line is the regression line for all genes. Genes enriched in secondary siRNAs in ADAR mutant worms are marked in cyan (A,B) and their regression line is presented in cyan. Genes enriched in primary siRNAs in ADAR mutant worms are marked in purple (A,B) and their regression line is presented in purple. 3'UTR edited genes are marked in cyan (C,D) their regression line is presented in cyan. *egl-2* and *lem-2* genes are marked in blue and yellow respectively in (C,D).

A.

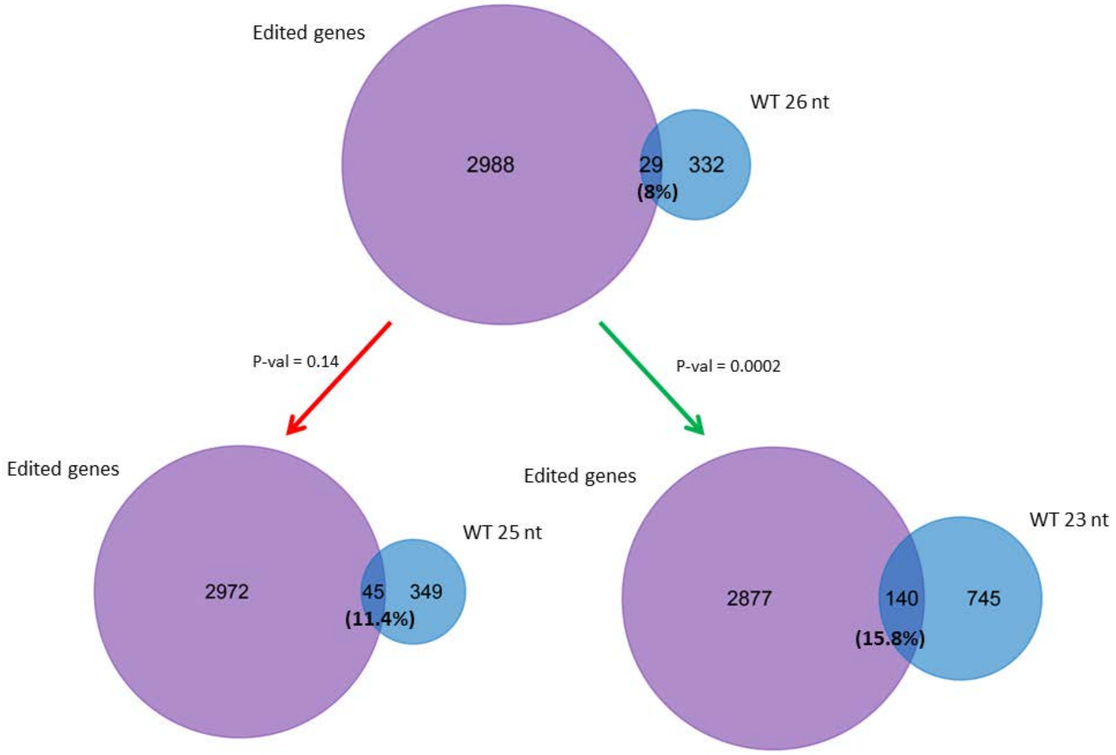

Edited genes
WT 26 nt
2988
29
332
(8%)
P-val = 0.14
P-val = 0.0002
Edited genes
WT 25 nt
2972
45
349
(11.4%)
Edited genes
WT 23 nt
2877
140
745
(15.8%)


B.

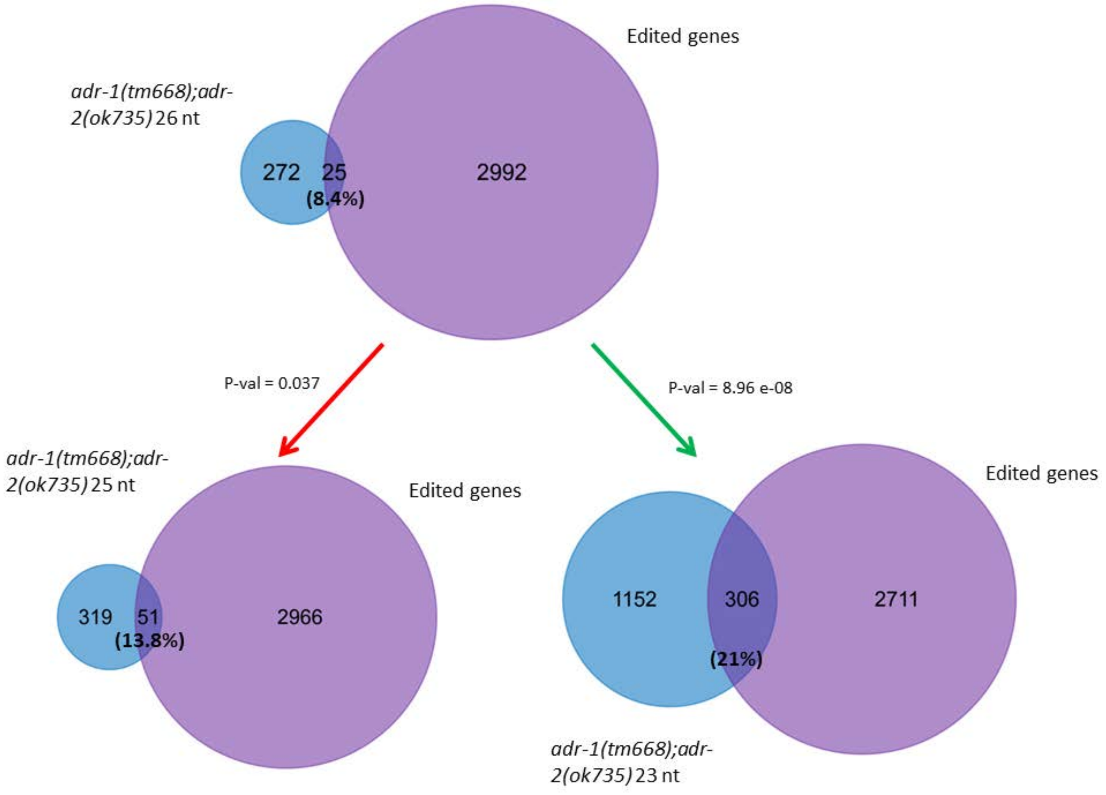

Edited genes
*adr-1(tm668);adr-2(ok735)* 26 nt
272
25
(8.4%)
2992
P-val = 0.037
P-val = 8.96 e-08
*adr-1(tm668);adr-2(ok735)* 25 nt
Edited genes
319
51
(13.8%)
2966
Edited genes
1152
306
2711
(21%)
*adr-1(tm668);adr-2(ok735)* 23 nt

C.

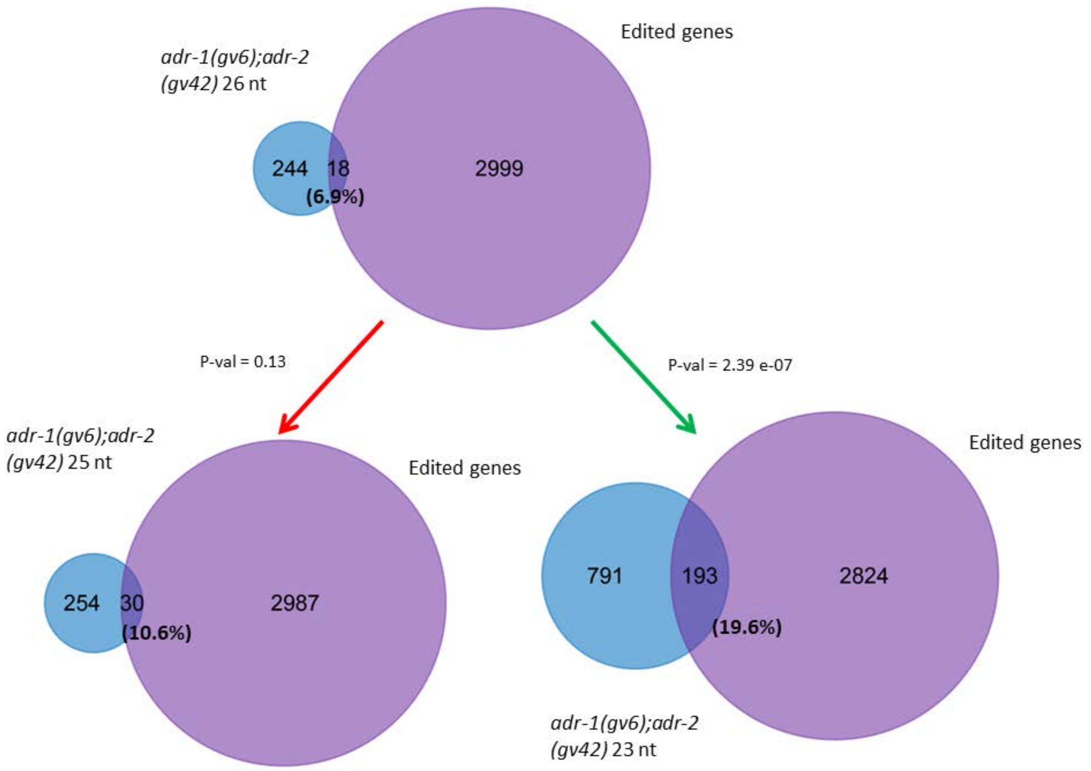


**Supplementary Figure 2. 23nt long antisense primary siRNAs are more aligned to edited genes than 25nt long or 26nt long siRNAs.** Venn diagrams showing overlap between genes that have primary antisense siRNAs with different sizes (23nt, 25nt, 26nt) aligned to them and edited genes. A. overlap in wildtype worms, B. Overlap in *adr-1(tm668);adr-2(ok735)* mutant worms, and C. overlap in *adr-1(gv6);adr-2(gv42)* mutant worms. We calculated if there is a significant difference in the overlap between the number of edited genes with antisense primary siRNAs alignment that are 26nt long to 25nt long and 23nt long by using Fisher exact test. Significant comparison is marked with P-value and green arrow, and non-significant comparison is marked by P-value and red arrow. As can be seen, in all strains, there is a statistically significant increase in the correlation of siRNA expressed genes to edited genes when comparing 23nt expressed genes to 26nt expressed genes. The difference in correlation to edited genes between 25nt expressed genes and 26nt expressed genes was not significant for any strain.

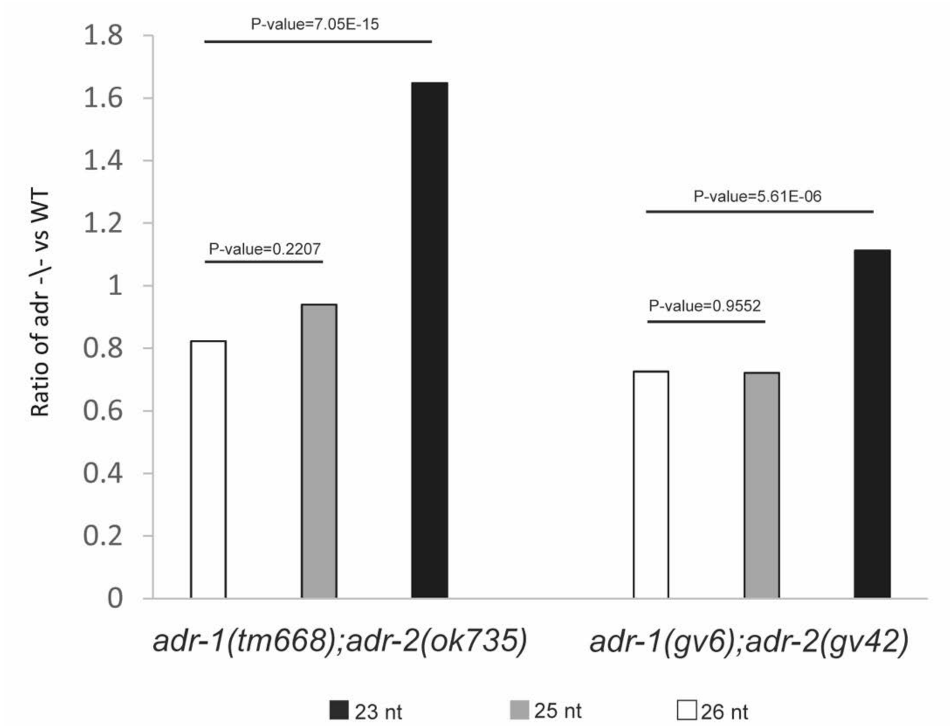


**Supplementary Figure 3. 23nt long antisense siRNAs aligned to edited genes are significantly enhanced in ADAR mutants compared to Wildtype worms.** Bar graphs showing the ratio of the number of primary antisense siRNA aligned to edited genes of different sizes between N2 worms and *adr-1(tm668);adr-2(ok735)* mutant (left) and *adr-1(gv6);adr-2(gv42)* mutant (right). 26nt long siRNAs are in white, 25nt long siRNAs are in gray, 23nt nt long are in black. Comparison between the sizes was done by Fisher exact test (p-values are marked in the graph).

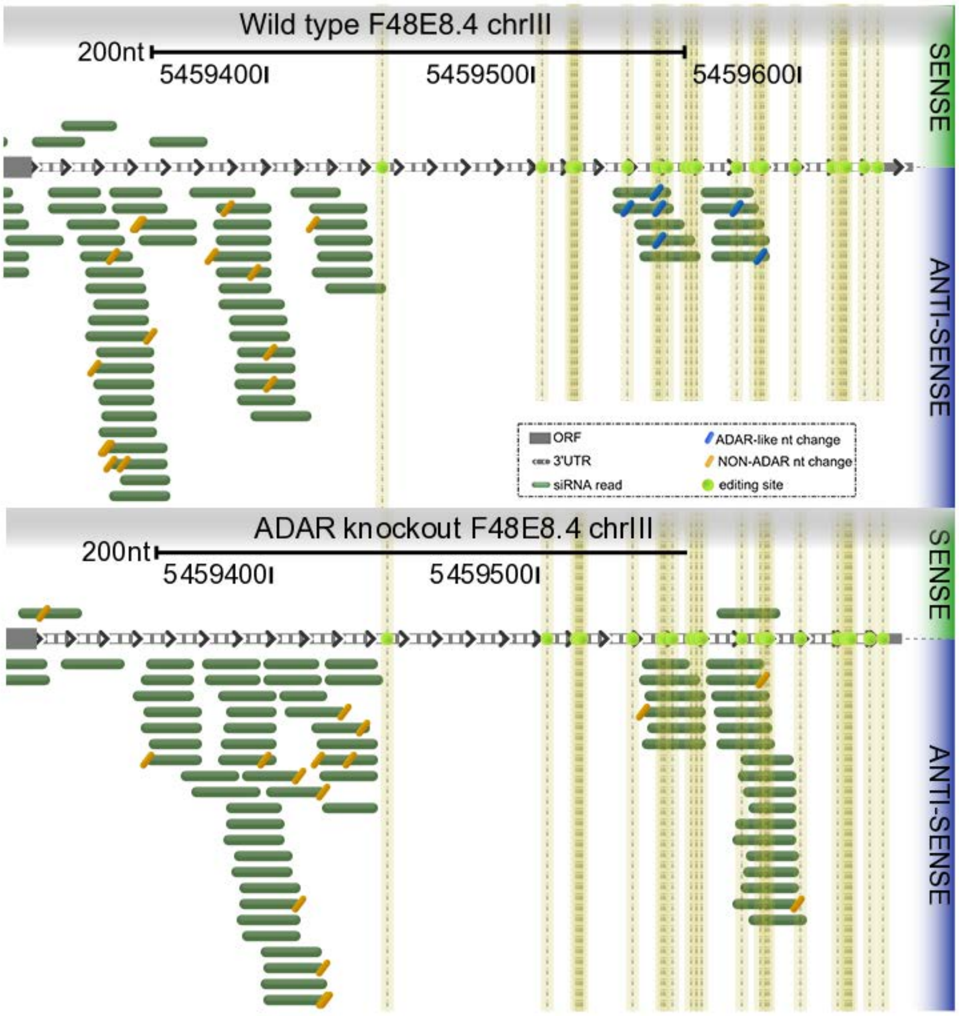


**Supplementary Figure 4. Example of secondary siRNAs aligned to an edited gene.** Presentation of secondary siRNAs sequences (green) aligned to the 3'UTR of gene F48E8.4 in wildtype and ADAR mutants. Editing sites are marked by yellow lines. Only secondary siRNAs that originated from wildtype worms contain the editing change (T-to-C, in blue).

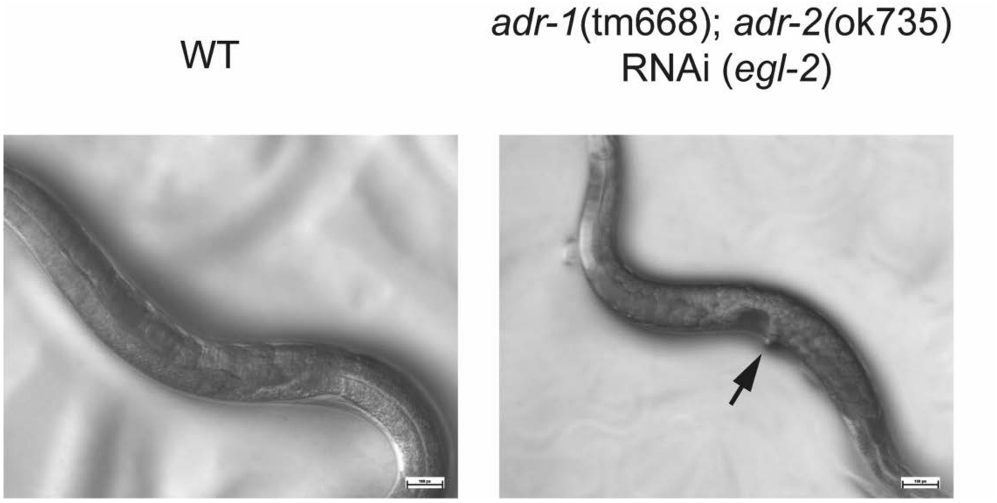


**Supplementary Figure 5. *egl-2* mutant knockdown morphogenesis defects.** Images of wildtype (left), and *adr-1(tm668) I; adr-2(ok735) III* that underwent RNAi targeted to *egl-2* worms (right) are shown. The arrow indicates one of the *egl-2* mutant phenotypes, the protruding vulva phenotype (pvl). Wildtype adult worm with normal body morphology is shown. Scale bar: 100 µm.

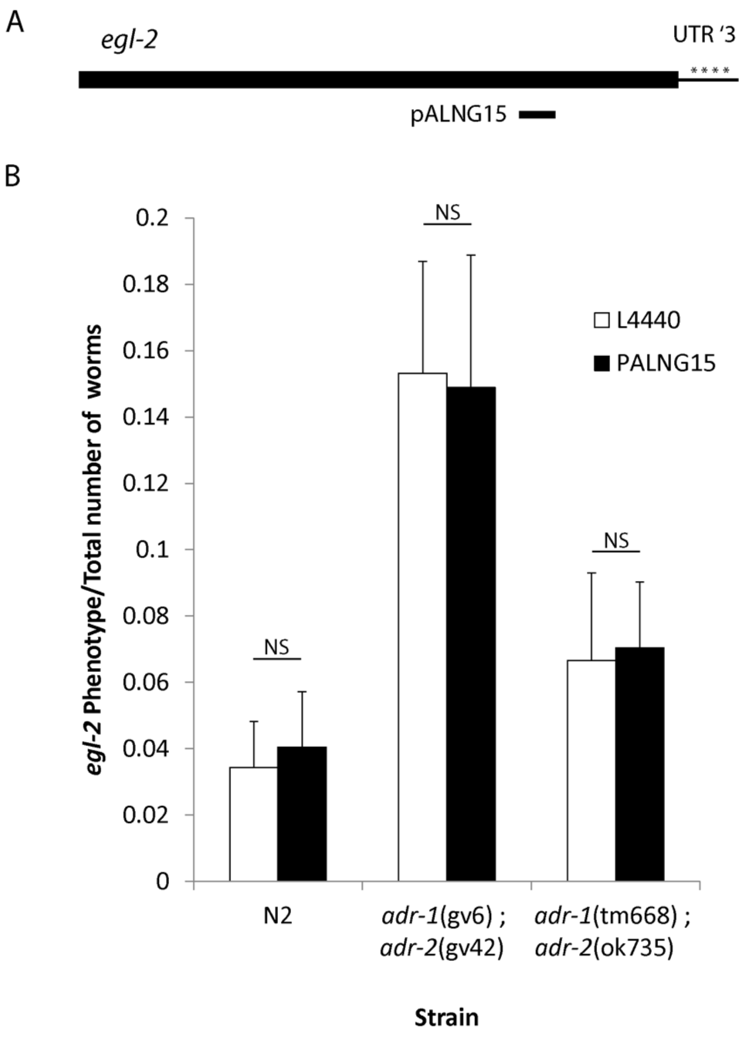


**Supplementary Figure 6. RNAi triggered against non-edited region in *egl-2* gene does not yield phenotypes in ADAR mutant worms.** Quantification of the fraction of *egl-2* phenotype from total worms observed in the RNAi experiments. PALNG015 is an RNAi plasmid against exon 2 in *egl-2*, which does not undergo editing. L4440 is a control RNAi vector. Strains that were tested are N2, wildtype worms, BB4, *adr-1(gv6);adr-2(gv42)* mutant worms, and BB21 *adr-1(tm668);adr-2(ok735)* mutant worms. Abnormal phenotypes that were scored include bloated worms, exploded worms, and bag of worms phenotypes. P-value was calculated from at least 3 biological replicates. NS for non-significant.

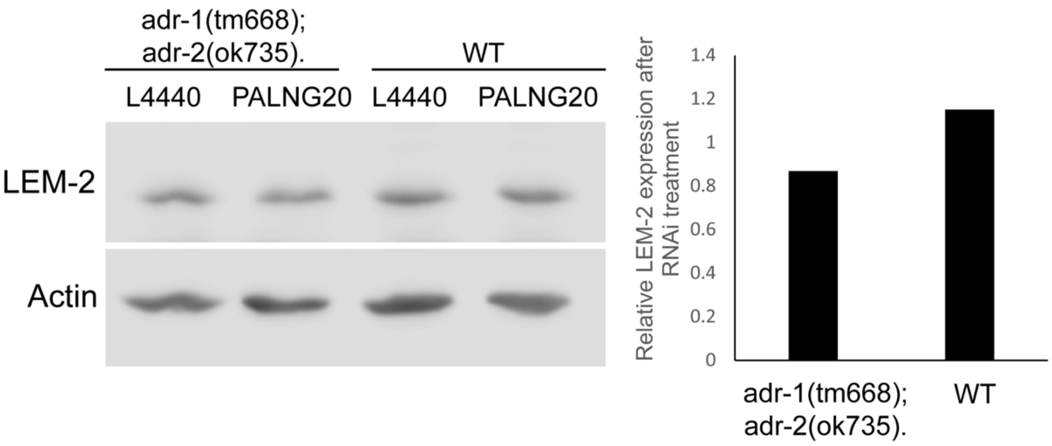


**Supplementary Figure 7. An example of a western blot quantifying LEM-2 protein level after RNAi triggered against the edited region in its 3'UTR.** Western blot analysis of the level of LEM-2 protein in wildtype (WT) worms and ADAR mutant (*adr-1(tm668); adr-2(ok735)*) worms treated with empty vector (L4440) or *lem-2* 3'UTR RNAi (PALNG20). Actin was used for loading control and background quantification. The bar graph shows the LEM-2 levels normalized to the Actin and relative to the empty vector (L4440). Single replicate blot is shown from several biological repeats, summarized in Figure 4.

C54D1.5 genomic DNA
chrX: 7144714-7144741
Position:7144729

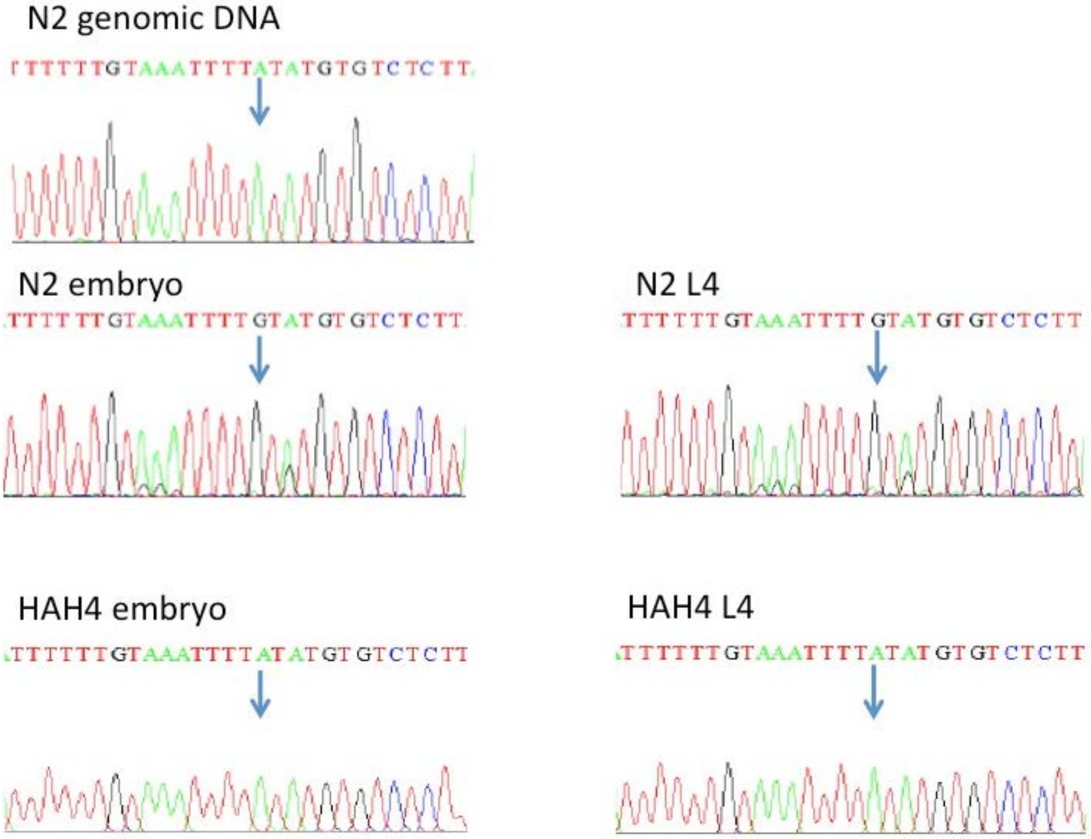

***F48E8.4***

chrIII: 5459498…5459554

Positions: 5459503, 5459514, 5459515, 5459516, 5459535, 5459546, 5459547, 5459550

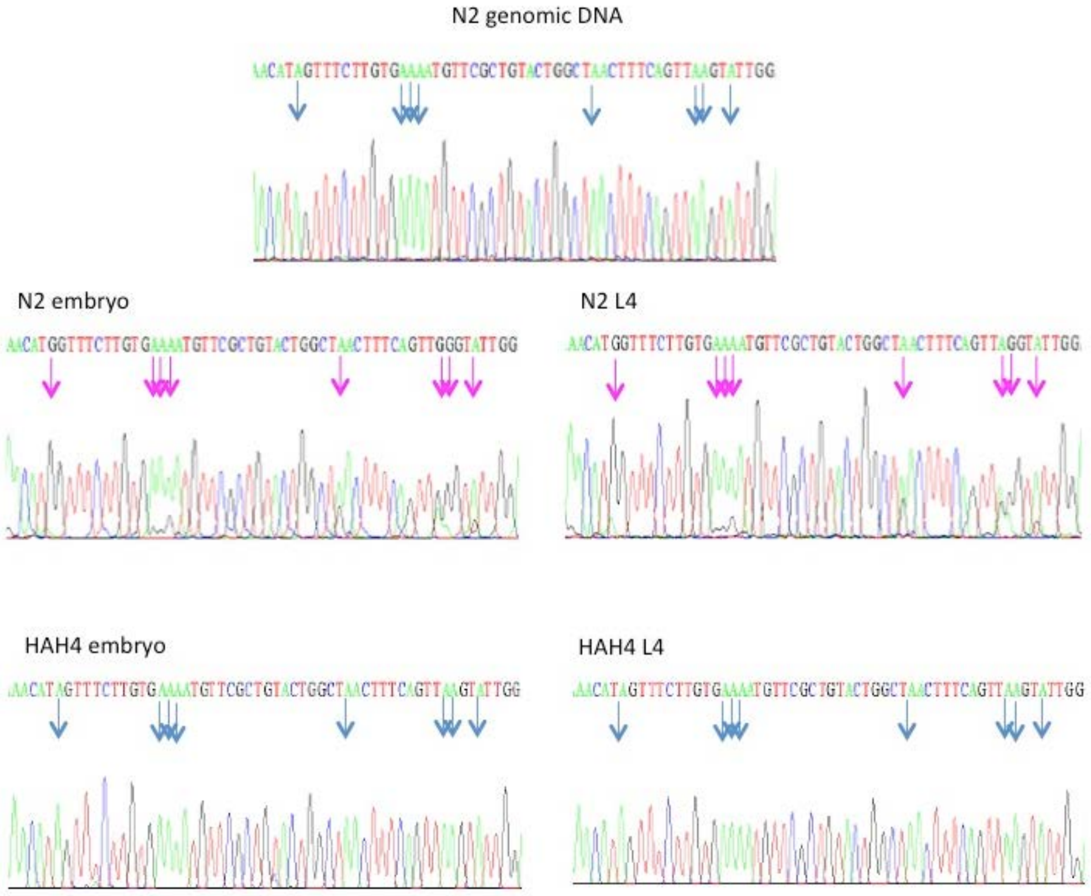

### *egl-2* genomic DNA

chrV: 1289532…1289444

Positions: 1289524, 1289508, 1289496, 1289485,1289468, 1289447

#### N2 genomic DNA

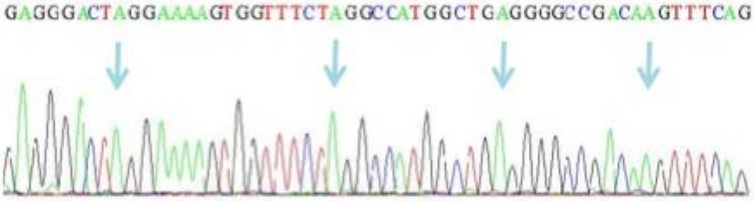


#### N2 embryo

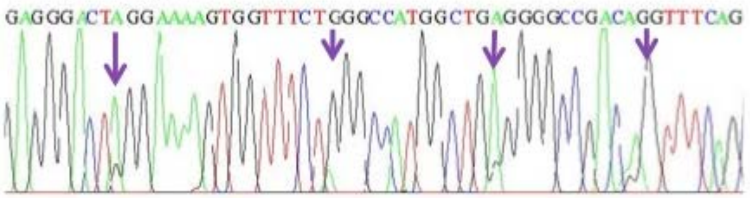


### N2 L4

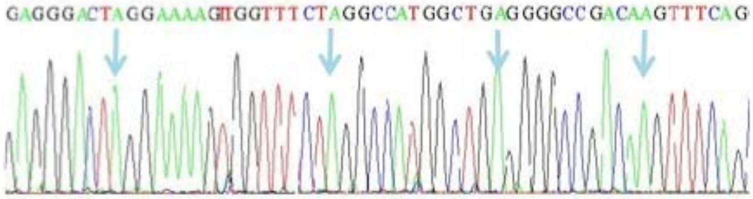


#### HAH4 embryo

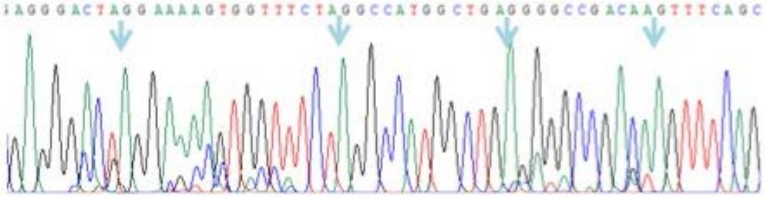


#### HAH4 L4

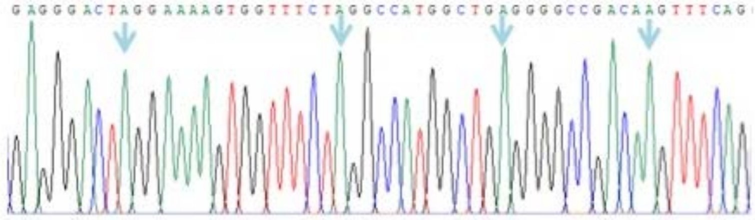

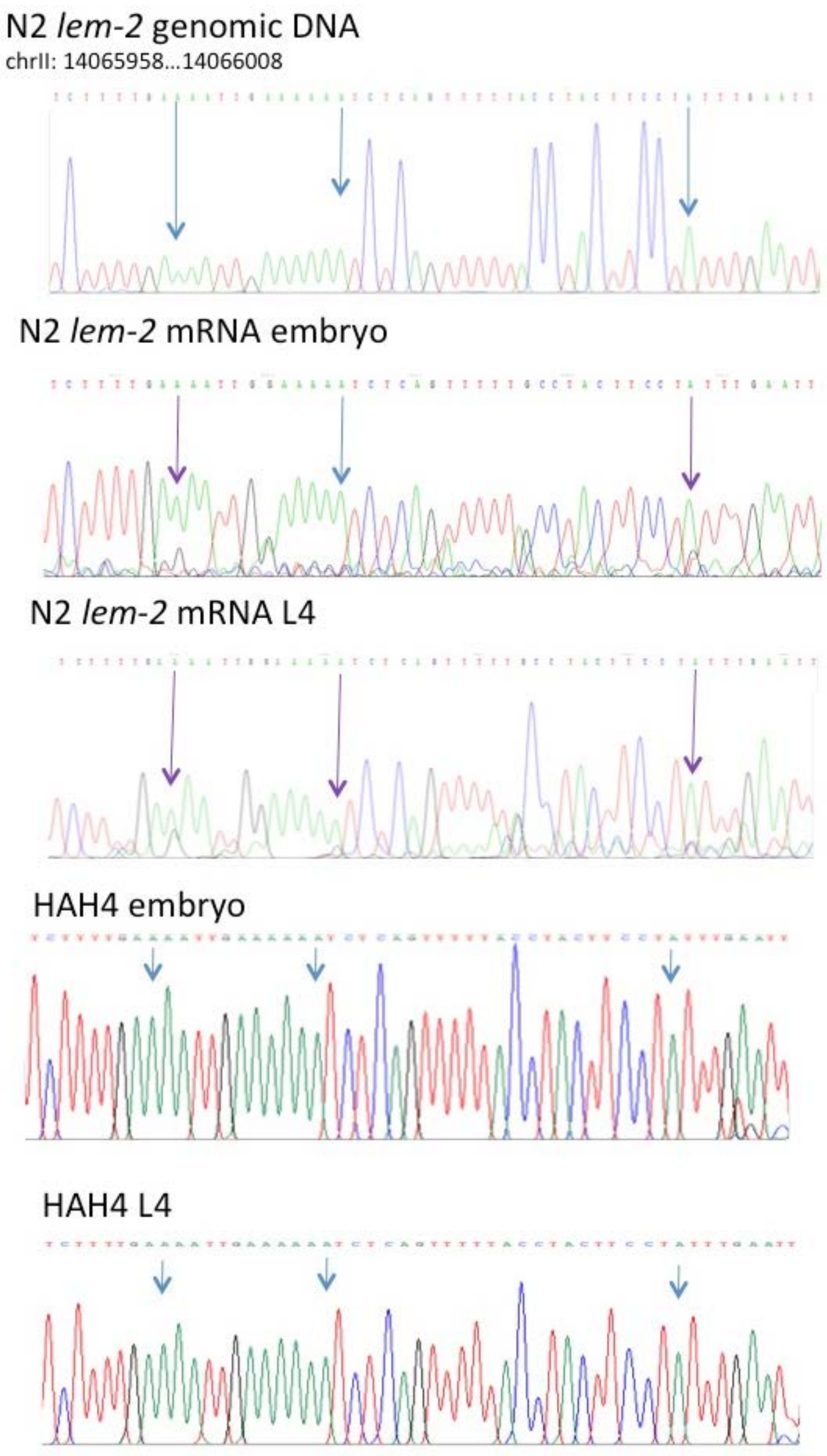


**Supplementary Figure 8. Genes are not edited in HAH4 strain *adr-2 (G184R)* mutant**. Sanger sequencing of *C54D1.5*, *F48E8.4* , *egl-2*, and *lem-2* genomic DNA and mRNA from embryo stage and from L4 stage from N2 or HAH4 (*adr-2(G184R))* strains. Blue arrows indicate that no editing was observed, magenta arrows indicate edited site.

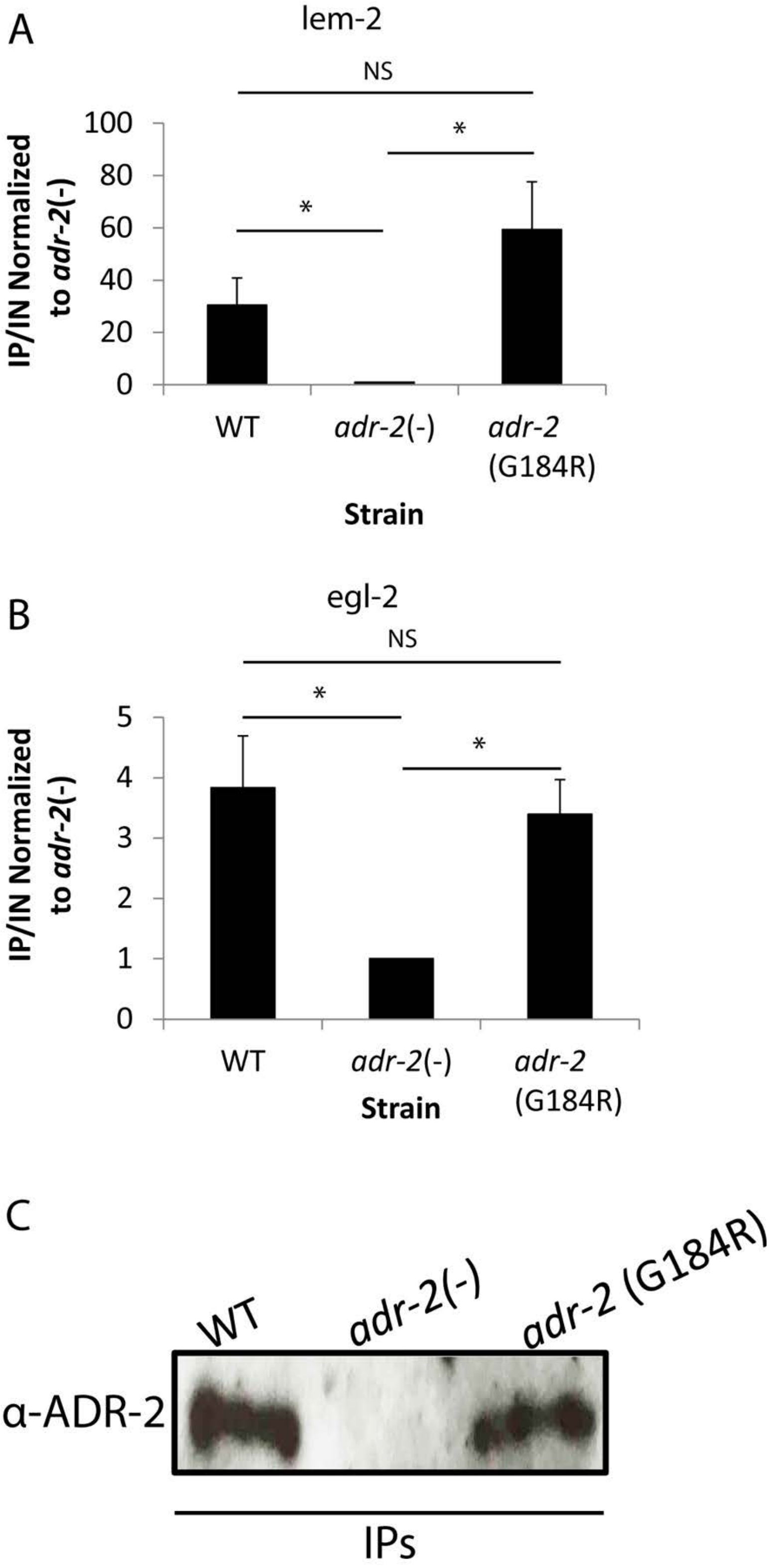


**Supplementary Figure 9. No significant changes in *egl-2* and *lem-2* RNA binding ability by ADR-2 in the WT and *adr-2 (G184R)* Strains.** (A-B) qPCR quantification of ADR-2 protein binding to *egl-2* (A) or *lem-2* (B) RNA by α-ADR-2 RIP on Wildtype (WT), *adr-2 (ok735)* (presented as adr-2 (-)) and *adr-2 (G184R)* worms. Data include 3 biological replicates. * P-value <0.05; NS for non-significant P-value. (C) Western blot sample of one biological replicate of worm lysate from strains wildtype (WT), *adr-2(ok735)* (presented as adr-2(-)) and *adr-2(G184R)* used for the RNA Immunoprecipitation qPCR (RIP- qPCR). The western blot shows that the antibody detects ADR-2 protein specifically and that the deamination mutation in ADR-2 *adr-2(G184R)* produces protein.

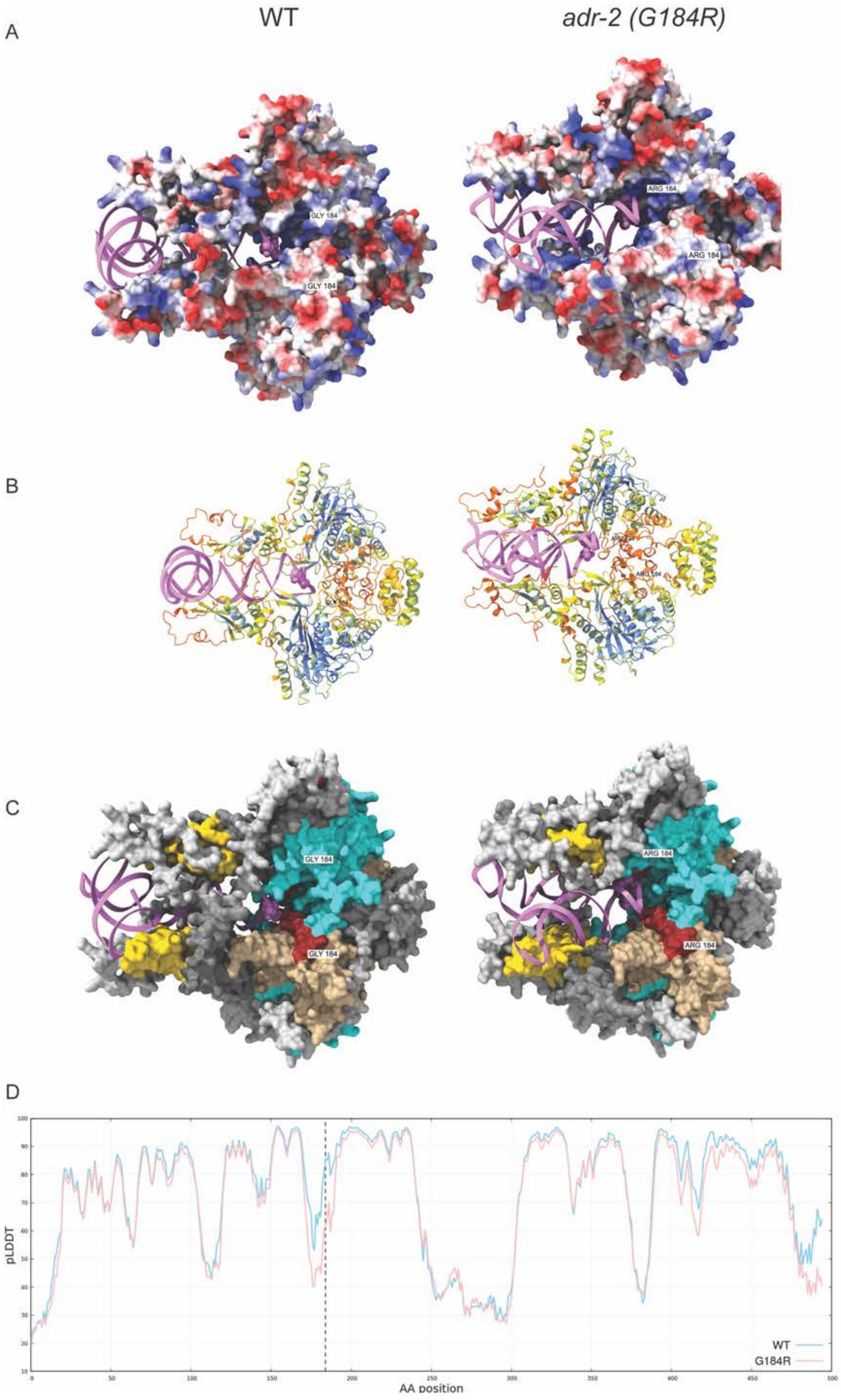

WT
adr-2 (G184R)
A
GLY 184
GLY 184
ARG 184
ARG 184
B
C
GLY 184
GLY 184
ARG 184
ARG 184
D
pLDDT
AA position
WT
G184R

**Supplementary Figure 10. The structural model of ADR-2; ADBP-1 and RNA shows a significant change in RNA binding when ADR-2 is mutated.** (A-C) Tetrameric model of ADR-2X2 and ADBP-1X2 from (Mu et al., 2025) with *egl-2* RNA using AlphaFold3. On the left is ADR-2 wildtype and on the right ADR-2 with G184R mutation in pink is *egl-2* RNA. Spheres in the RNA are editing sites. A. Color coded by the electrostatic potential. Blue represents positive potential, and red represents negative potential. B. Ribbon representation color coded by plDDT. C. Similar to Figure 6C. Areas of higher flexibility because of the G184R mutation are marked (see the positions in D). In brown are aa positions 160-190. In tan are aa positions 400-495. D. Plot of pLDDT values per residue for the WT (blue) and G184R (red) obtained from the AlphaFold3 models. The dashed line represents the position of the mutation.

### Supplementary Tables

**Supplementary Table 1. Expression changes in mRNA, primary or secondary siRNAs between wildtype and ADAR mutants.**

**Supplementary Table 2. pyDock energy terms**

| ADR-2 WT/G184R | Model number | Ele | Desolv | VDW * 0.1 | Total |
|---|---|---|---|---|---|
| WT | 1 | -316.560 | 135.921 | -5.6 | -186.249 |
| G184R | 1 | -383.725 | 150.160 | -2.4 | -236.043 |
| WT | 2 | -329.975 | 138.686 | 7.6 | -183.676 |
| G184R | 2 | -336.295 | 126.092 | -2.0 | -212.280 |
